# A genome wide association study identifies a new variant associated with word reading fluency in Chinese children

**DOI:** 10.1101/2022.10.23.513381

**Authors:** Zhengjun Wang, Shunan Zhao, Liming Zhang, Qing Yang, Chen Cheng, Ning Ding, Zijian Zhu, Hua Shu, Chunyu Liu, Jingjing Zhao

## Abstract

Reading disability exhibited defects in different cognitive domains, including word reading fluency, word reading accuracy, phonological awareness, rapid automatized naming, and morphological awareness. To identify the genetic basis of Chinese reading disability, we conducted a genome wide association study (GWAS) of the cognitive traits related to Chinese reading disability in 2284 unrelated Chinese children. Among the traits analyzed in the present GWAS, we detected one genome wide significant association (*p*<5×10-8) on reading fluency for one SNP on 4p16.2, within EVC genes (rs6446395, *p*=7.55×10^−10^). Rs6446395 also showed significant association with word reading accuracy (*p*=3.39×10^−4^), phonological awareness (*p*=7.12×10^−3^), and rapid automatized naming (*p*=4.71×10^−3^), implying multiple effects of this variant. Gene-based analyses identified a gene to be associated with reading fluency at the genome-wide level. The eQTL data showed that rs6446395 affected EVC expression in the cerebellum. Our study discovered a new candidate susceptibility variant for reading ability and provide new insights into the genetics of development dyslexia in Chinese Children.

## Introduction

Developmental Dyslexia (DD, also known as reading disability) is a complex genetically influenced neurodevelopmental disorder characterized by difficulties in acquiring fluent reading skills despite normal intelligence and adequate education[1]. It is known to be a hereditary neurological disorder with a prevalence of approximate 1.3 - 17.5 % among children varying in different orthographies[2,3]. The genetic heritability of DD and reading-related skills were high (40%-80%)[4]. Specifically, the heritability estiamtes of reading accuracy and reading fluency were 0.03-0.60[5] and 0.20-0.73[6,7], respectively.

A few studies have explored the genetic basis of dyslexia, using either candidate genetic analysis or genome-wide association studies (GWAS). Several candidate loci for dyslexia have been identified through linkage mapping, including DCDC2, KIAA0319, DYX1C, and MRPL19, although variants in those genes and loci were not replicated in different populations[8-11]. Meanwhile, in the last decades, a few GWAS have identified some genetic association with dyslexia, but few of those associations have reached genome-wide significance and were not replicated in independent datasets either[12-16]. It should be noted that a recent GWAS study identified 42 genome-wide significant loci associated with dyslexia in 51,800 adults self-reporting a dyslexia diagnosis and 1,087,070 controls, and validated 23 loci in independent cohorts of Chinese and European ancestry[17].

Other than GWAS with dyslexia, more and more candidate gene studies and GWAS started to investigate genetic correlates with quantitative trait of reading ability/disability and reading/dyslexia-related cognitive predictors in recent years[18-20]. Reading accuracy and reading fluency are two important aspects to account reading ability/disability. Previous GWAS on reading ability have focused more on word reading accuracy but have largely ignored word reading fluency[14-16,21]. This study for the first time investigated the genetic associations of word reading fluency using GWAS. Reading fluency refers to a reader’s ability to read quickly and accurately[21-23]. Previous research findings have consistently shown that impaired reading fluency is one of the core deficits of dyslexia[23,24]. Especially for older dyslexic children, slow reading rate is a major difficulty since they have to read connected text[25]. In addition, dyslexic readers may perform poorly in reading comprehension due to low reading fluency (impaired decoding skills and/or slow reading rate)[26,27]. It is generally agreed that reading fluency can be used for diagnosis of DD in transparent languages, while reading accuracy is the main criteria for defining DD in languages with deep orthography.

Although dyslexia is typically characterized by genetically influenced impairment in word reading accuracy and word reading fluency, it is generally agreed that genetic influence engenders cognitive disabilities, which in turn affect reading acquisition and lead to dyslexia. The well-accepted phonological deficit theory suggests that developmental dyslexia is mainly caused by deficits in three dimensions of phonological skills: phonological awareness (PA, e.g., phoneme deletion), phonological processing speed (e.g., rapid automatized naming (RAN) of objects and digits), and phonological representation (e.g., digit span for verbal short-term memory (VSTM))[28,29]. Among which, RAN and PA played major roles while digit span played minor roles[30]. Moreover, morphological awareness (MA) has also been found strongly correlated with reading ability[31]. PA, RAN, digit span, MA therefore might be useful endophenotypes for research of susceptibility genes in dyslexia.

Indeed, recent GWAS attempted to examine cognitive endophenotypes to study the genetics of reading ability, which may share, in part, the underlying genetic influences of dyslexia. Gialluisi and colleges conducted a genome-wide association study with cognitive traits of PA, RAN, and digit span [32]. They observed a genome-wide significant association (rs17663182, on 18q12.2 within MIR924HG) on rapid automatized naming of letters in the European populations (N = 2562–3468).

Almost at the same time, genome-wide significant association of another loci, rs1555839 (10q23.31, upstream of PRL7P34), were observed with RAN and rapid alternating stimulus (RAS), in Hispanic American and African American youth (n=1331, p<5*10-8). This rs1555839-RAN association was replicated in an independent cohort from Colorado[32].

Up to date, all previous GWAS on reading ability/disability and reading/dyslexia related cognitive traits were mainly performed in the Western population whose genetic backgrounds are different from the Chinese population. Although dyslexic candidate genes, such as KIAA0319, DCDC2, and DYX1C1, have been shown to be associated with dyslexia in Chinese children[33-36], no GWAS of reading ability and relative cognitive traits had been conducted in Chinese children until now. In fact, so far there is still no well-recognized definition criteria for Chinese developmental dyslexia. The cognitive deficits of Chinese developmental dyslexia is also in debate. Recently, our lab used reading accuracy as the main criteria for defining Chinese developmental dyslexia and found the prevalence of Chinese dyslexia is around 10% in primary school children[37]. We also observed in this study that about 75% of Chinese developmental dyslexia can be accounted for phonological deficits, combining both phonological awareness deficit and rapid automatized naming deficit[37]. Moreover, in vast contrast to the alphabetic languages, as words in Chinese written language are composed with different morphemes[38,39], morphological awareness has also been identified as one of the core deficits in Chinese developmental dyslexia[40].

To understand the genetic basis of reading ability and related cognitive traits, we conducted GWAS in 2284 Chinese children. We analyzed eight behavioral and cognitive traits in this Chinese cohort, including word reading accuracy, word reading fluency, phoneme awareness, RAN (digit, picture, color, and dice), and morphological awareness. We identified a genome-wide significant association at 4p16.2 (rs6446395) with word reading fluency. This genetic variant was also correlated with word reading accuracy and phonological skills. Gene-level analyses presented a significant enrichment of a gene associated with reading fluency. Furthermore, a series of bioinformatics analyses were performed to investigate the function of the associated loci. The findings may provide new evidence for the genetic etiology of dyslexia and related cognitive traits.

## Material and methods

### Participants

A total of 3127 primary students aged 9 to 14 years from three cities and four districts in China (Xi’an-YT, Xi’an-CB, Qingyang, and Baotou) were recruited in this study. They were all school-age children with normal intelligence, according to the description of parents and teachers. Finally, 2480 participants were eligible for subsequent genotyping and association analysis. The sample size for GWAS of behavioral and cognitive traits are as follows: word reading accuracy (N = 2270), word reading fluency (N = 2270), phoneme awareness (N = 2244), rapid automatized naming (N = from 1968 to 2080), and morphological awareness (N =2076). Ethical approval was obtained by Ethics Committee of Shaanxi Normal University and written informed consent was obtained for all the participant children’s parents.

### Phenotypic measures

We performed GWAS on eight phenotypes: word reading accuracy (namely character recognition, CR), word reading fluency (RF), phoneme awareness (PA), rapid automatized naming (RAN) of digit, picture, color, and dice, and morphological awareness (MA). The details explanations of the phenotypic measures were described in Supplementary Methods. These traits showed moderate to high correlation between traits (Table S1).

### Genotype quality control and imputation

DNA was extracted from saliva samples of 2477 participants, and individuals were genotyped using Illumina Asian screening array (650K) by Beijing Compass Biotechnology. Genotype quality control was performed as previously described in Plink v1.90[41]. Briefly, SNPs were filtered out if they showed a variant call rate <0.95, a minor allele frequency (MAF) <0.05, a missing genotype data (mind) <0.90 (4 samples were removed), or a hardy-weinberg equilibrium (HWE) <10^−5^ with each dataset. Samples showing mismatches between genetic and reported sex (8 samples), or unexpected duplicates or probable relatives (PI-HAT>0.20, 53 samples) were discarded[42,43]. Additional 128 samples were discarded due to missing phenotypes. After quality control, 2284 children with 411091 SNPs were left.

For imputation, autosomal variants were aligned to the 1000G genomes phase 1v3 reference panel. Imputation was performed using Michigan imputation Server 4.0 in 5Mb chunks with 500kb buffers, filtering out variants that were monomorphic in the Genome Asia Pilot (GAsP). Chunks with 51% genotyped variants or concordance rate <0.92 were fused with neighboring chunks and re-imputed. Finally, imputed variants were filtered out for rsq <0.60, MAF<0.05, mind<0.1, HWE<10^−5^ using Plink (v1.90). After quality control, 2284 children with 4173723 SNPs were included for the final genome-wide association analyses.

### Genome-wide association analyses

After quality control and imputation, genome-wide association analyses were performed using Plink, fitting an additive model to the linear regression model with adjustment for sex, age, and the first five principle components of population structure[42]. A Manhattan plot of -log^10^P was generated using the ggplot2 package in R. SNPs with *p* < 5×10^−8^ was considered genome-wide significance.

### Assessment of SNPs previously associated with dyslexia and related traits

The candidate SNPs in 5 genes (DCDC2, DYX1C1, ROBO1, KIAA0319, and MPRL19) previously reported in DD were selected for replication, including 27 SNPs (rs2760157, rs12193738, rs3903801, rs807507, rs16889506, rs3212236, rs761100, rs6732511, rs4504469, rs3756821, rs2179515, rs2038139, rs1000585, rs2038137, rs793862, rs3743204, rs807701, rs807724, rs917235, rs1091031, rs16889556, rs699463, rs600753, rs6935076, rs9366577, rs714939, rs2143340) [44,45]. The variant of rs2038139 with a MAF less than 0.05 was removed from the replication analysis. Among the remaining SNPs, we selected the leading 16 SNPs (rs3756821, rs761100, rs807507, rs1000585, rs3743204, rs807724, rs807701, rs2143340, rs16889506, rs699463, rs9366577, rs793862, rs714939, rs1091031, rs6732511, rs2760157) based on linkage disequilibrium (r^2^ > 0.4) and validated these variants (see supplementary Table S2).

Moreover, variants associated with an important dyslexic related cognitive trait-rapid automated naming in previous GWAS were selected for replication[18,32], including 13 SNPs (rs6963842, rs11177505, rs4320486, rs17663182, rs17605546; rs34822091; rs16928927, rs4571421, rs76161559, rs4307051; rs200580547, rs17663182, rs17605546). However, only rs6963842 and rs4571421 were present in our data. None of other SNPs were genotyped or imputed in our data. Therefore, we also performed replication analysis for these two SNPs (rs6963842 and rs4571421) in this study.

### Gene- and pathway-based enrichment tests

Gene-based association analyses for the phenotypic traits were performed using MAGMA[46]. First, SNPs were assigned to protein-coding genes based on the position to the NCBI37.3. A total of 17184 genes (out of 17264 genes definitions read) included at least one SNP that passed internal QC, and were thus tested in gene-based enrichment analysis. Gene-based statistics were computed using the SNP association statistics calculated in the GWAS of phenotype. Given the number of genes, the Bonferroni-corrected genome-wide significance threshold for this analysis was set to *p*<0.05/17184=2.9×10^−6^.

Using the results of gene-based enrichments analysis, we carried out the pathway-based enrichments through a competitive gene-set analysis in MAGMA[47]. A total of 5497 canonical pathways were tested from the Molecular Signatures Database website (MSIGDB, c2.all.v7.0.entrez). To correct enrichments statistic for testing of multiple pathways, the significance threshold was 9.1×10^−6^. eMAGMA was also used to identify novel candidate genes of dyslexia based on expression regulation in 13 brain regions[47].

### Bioinformatics analysis

To explore the affected gene expression, the expression quantitative trait locus (eQTL) for rs6446395 was examined in the UK Brain Expression Cohort data set (GSE46706)[48] and Genotype-Tissue Expression (GTEx) Cohort data set. Detailed processing and exclusion criteria have been described previously, and eQTL analysis was described by Ramasamy et al[49]. The EVC-interacting genes were from PINA v2, and the EVC-coexpressed genes were from GeneMANIA[50, 51].

## Results

### Single-variant genome-wide associations

The demographic description, behavioral and cognitive phenotype data for the samples are present in Table 1. After quality control for the genotype data as well as phenotype and covariate data cleaning performed, a total of 2270 individuals tested with the reading ability (word reading accuracy and fluency) were analyzed in the GWAS study. For phoneme awareness, morphological awareness, and rapid automatized naming tasks, 1968-2244 samples were analyzed.

**Table 1.**
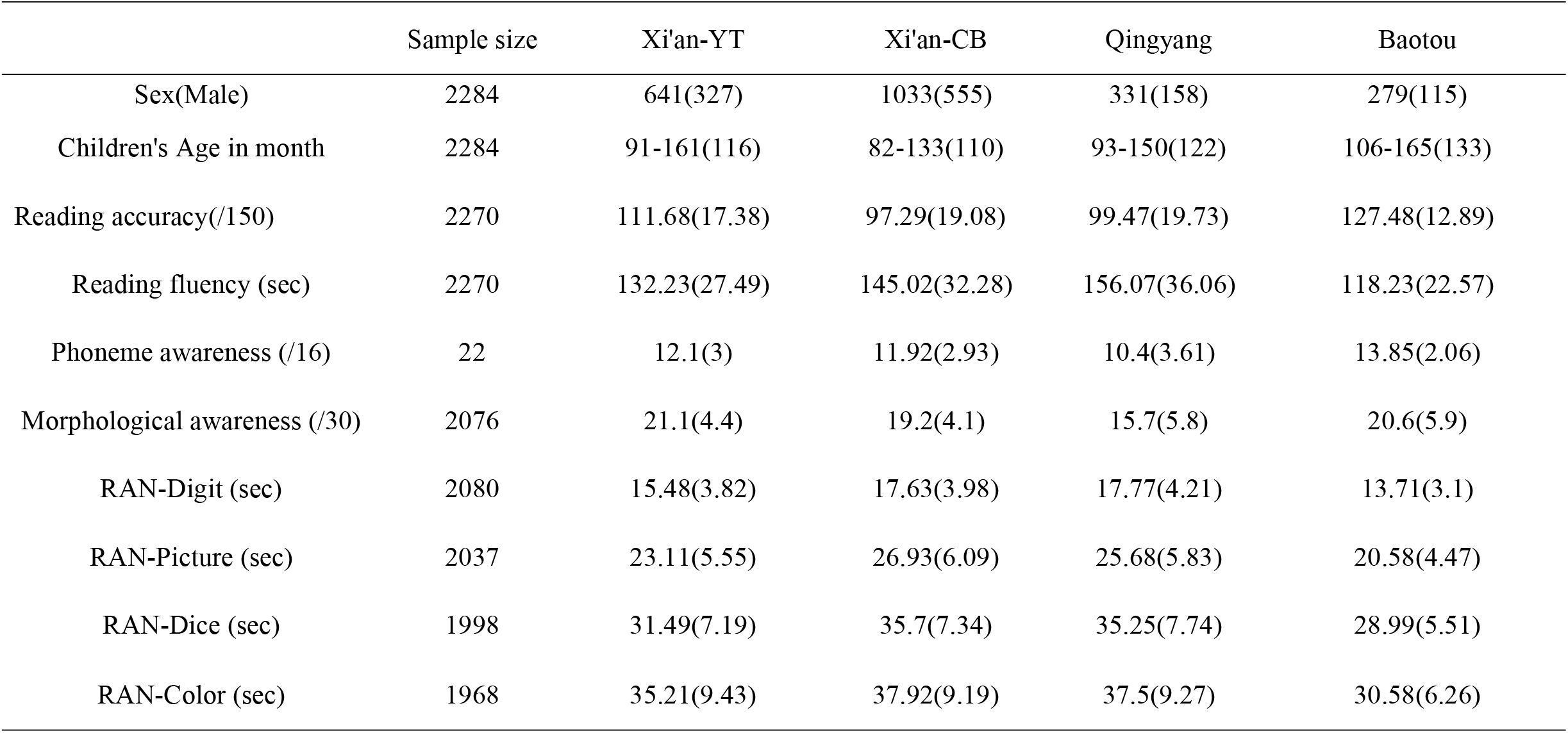
Main characteristics and recruitment criteria of the datasets involved in the present study.

Among the eight traits analyzed in the present GWAS, only reading fluency showed a variant of genome-wide significant association withstanding correction for multiple testing (*p*<5×10^−8^), mapped to chromosome 4p16.2. The significant association was observed for rs6446395 (G/A, MAF=0.442, *p*=7.33×10^−10^, major allele (G) β = −5.966, standardized β (SE)= −0.120(.020)). Rs6446395 was directly genotyped, mapped to the coding gene EVC (EvC ciliary complex subunit 1, Figure S1). Further details on the association of rs6446395 were reported in Figure 1. The variant rs6446395 showed consistent allelic trends for reading fluency in the different subsamples (Figure 2A).

**Figure 1.**
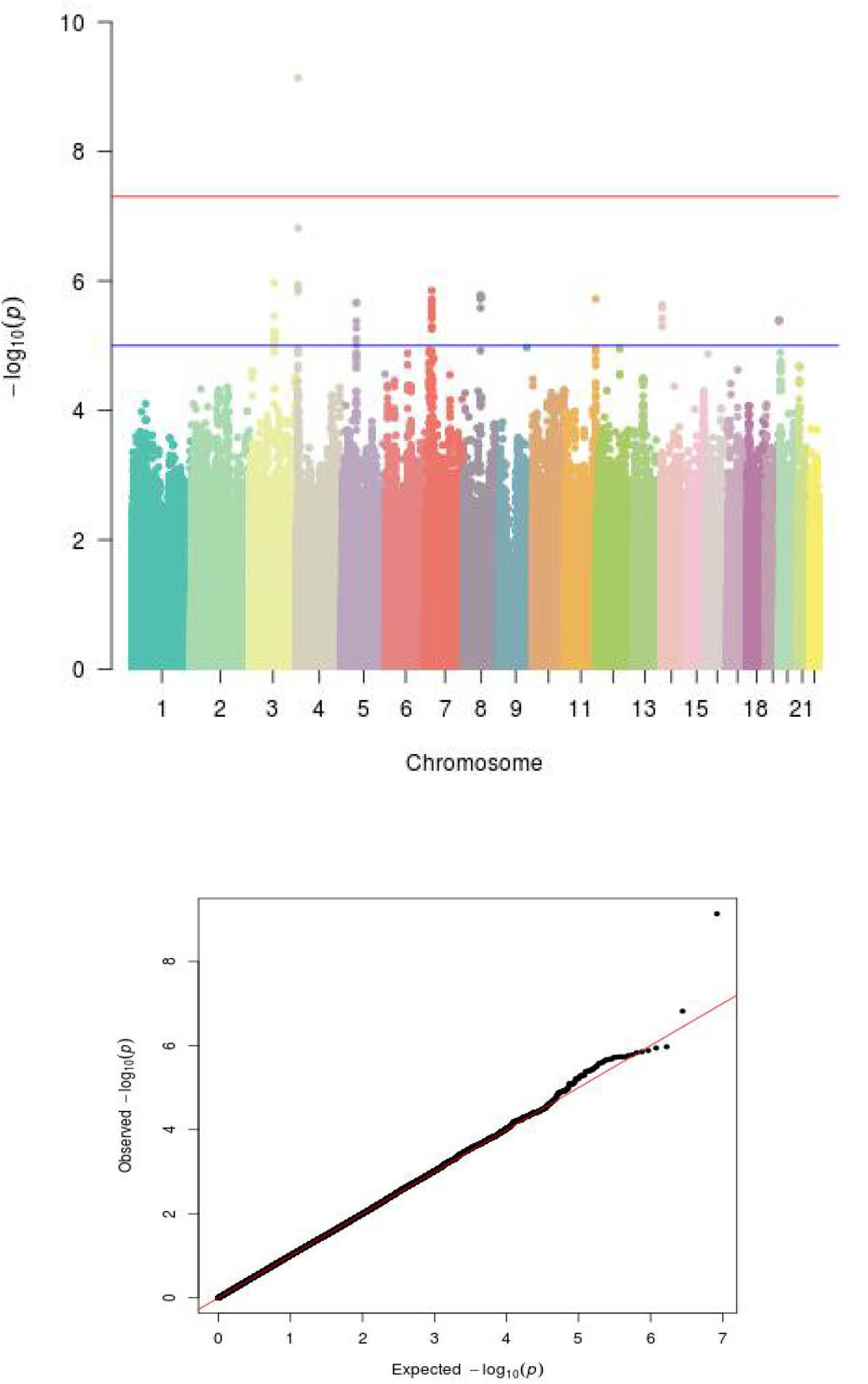
Manhattan plot summarising the results and QQ plots of the genome-wide association study (GWAS) for word reading fluency. The red line represent the genome wide significance threshold (*p*<5×10−8) λ= 1.017.

**Figure 2.**
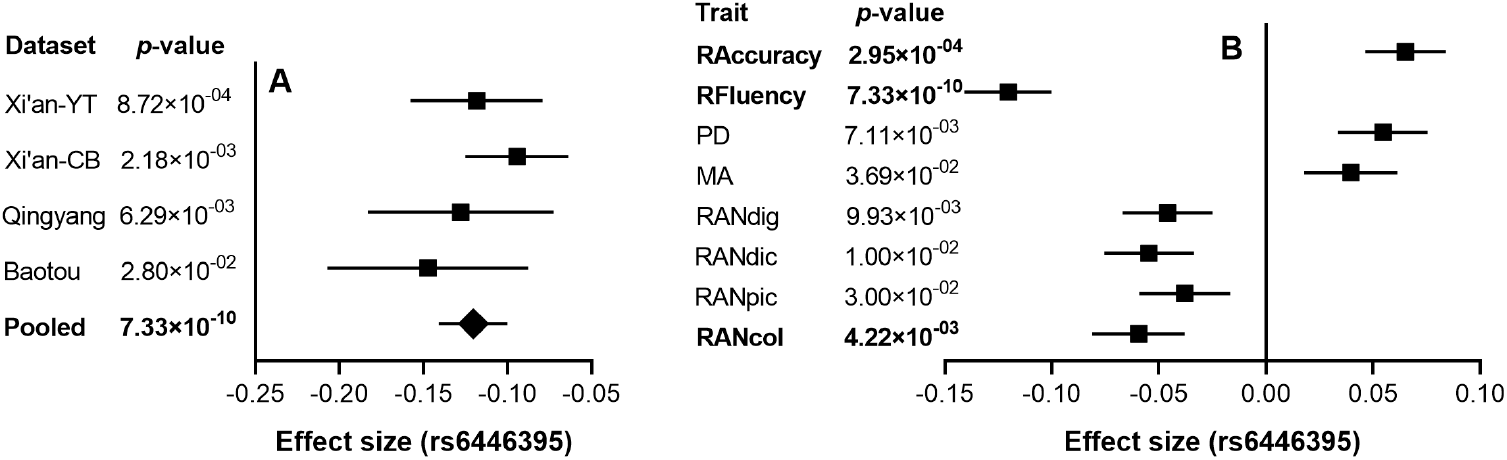
Forest plot of associations of rs6446395 with word reading fluency in populations from four districts (A) and with different cognitive traits (B) analyzed in the study. β refer to major alleles G.

Furthermore, rs6446395 showed weak association with character recognition (β=2.066, standardized β(SE)=0.065(0.019), *p*=2.95×10^−4^), phonological awareness(β=2.583, standardized β(SE)=0.055(0.021), *p*=7.11×10^−3^), and rapid automatic naming (digit: β=-0.327, standardized β(SE)=-0.046(0.021), *p*=9.93×10^−3^; picture: β=-0.483, standardized β(SE)=-0.038(0.021), *p*=3.00×10^−2^; color: β=-0.841, standardized β(SE)=-0.059(0.022), *p*=4.22×10^−3^; dice: β=-0.512, standardized β(SE)=-0.055(0.021), *p*=1.01×10^−2^;), and morphological awareness (β=0.349, standardized β(SE)=0.040(0.022), *p*=3.69×10^−2^) (see Figure 2B). The effects of genotypes (G/G, G/A, and A/A) in rs6446395 for word reading fluency and word reading accuracy were shown in Figure 3. Detailed results of GWAS analyses of other traits are reported in supplementary Table S5 and Figure S2-S8.

**Figure 3.**
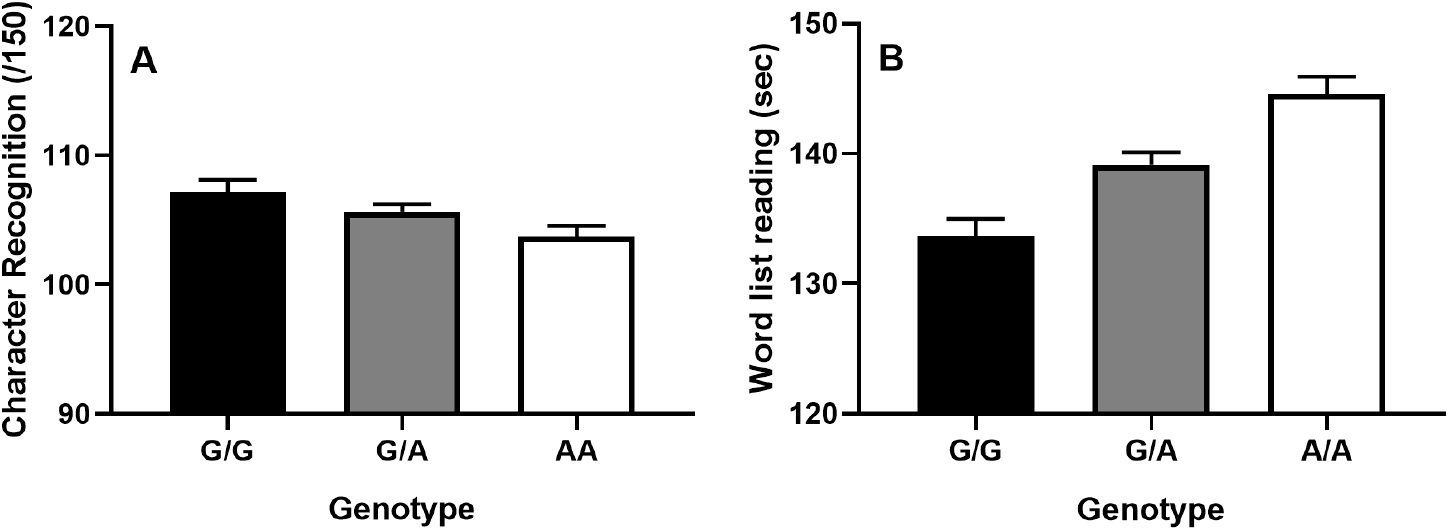
Boxplot of word reading fluency and word reading accuracy for the lead variant rs6446395. Genotype counts are G/G = 422; G/A = 1177; A/A = 685.

### SNPs previously associated with dyslexia and related cognitive trait

Among the 18variants selected for replication analysis, the most significant association were observed at rs2760157, an intronic SNP located in the KIAA0319 (6p22.2), with RAN-color (A/G, maf=0.438, *p*=6.97×10^−5^, major allele(A) β=1.103). Other variant, rs16889506, in KIAA0319 was also associated with RAN-color (C/T, maf=0.167, *p*=3.17×10^−4^, major allele(C) β = −1.391). Those associations showed the same direction of effects as in the original report. The details of the associations for all traits were shown in supplementary Table S2. No other SNP showed association that can survive test-wise correction (*p* > 0.0027=0.05/18).

### Genetic mechanism of the significant locus rs6446395

The significant locus (rs6446395) maps within the gene EVC. eQTL data from the UK Brain Expression Cohort showed that rs6446395 affected EVC expression in the cerebellar cortex (Figure 4). The minor allele G of rs6446395 is associated with decreased EVC expression in the cerebellum. EVC is expressed in brains at different development stages and throughout the human lifespan (see supplementary Figure S9). We found a nominally significant difference of expression of EVC in the cerebellum when participants had differnet genotypes of rs6446395 from GTEx Cohort (*p*=6.35×10^−2^). To further explore the specific functions of EVC, we mapped a network including EVC and its interaction or co-expressed genes. Six interacting genes and 20 co-expressed genes were observed indicating their association with neurocognitive function (see supplementary Figure S10).

**Figure 4.**
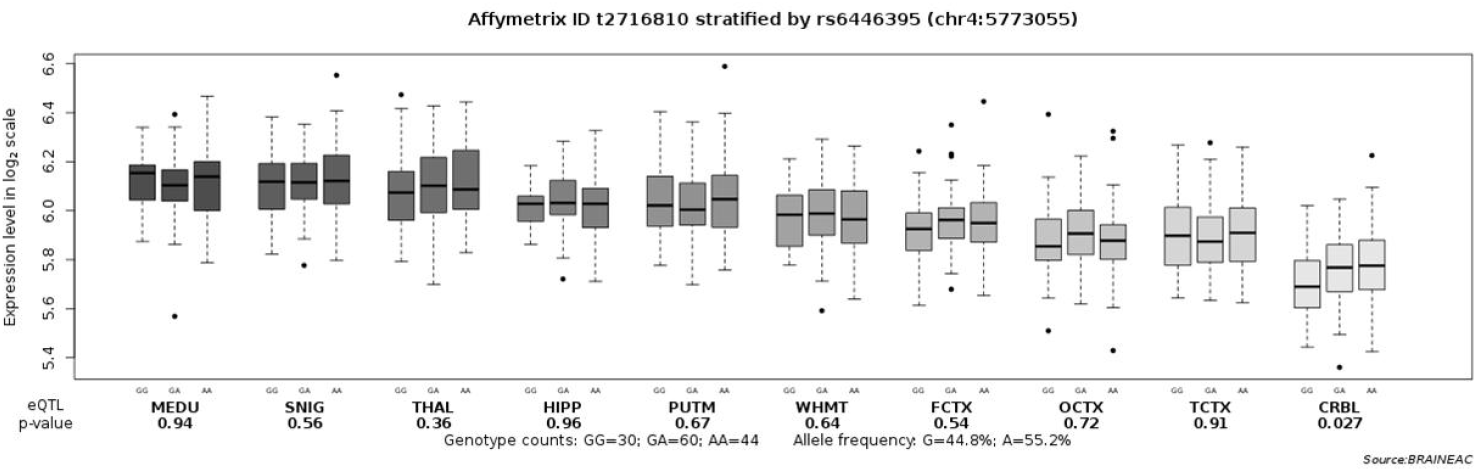
The effect of rs6446395 on EVC expression. The study of expression quantitative trait loci in brain tissue demonstrates the effect of rs6446395 on EVC gene expression in 10 different brain regions in 134 samples from the UK Brain Expression Cohort (UKBEC). Boxplot dashed bars mark the 25th and 75th percentiles. CRBL, cerebellar cortex; FCTX, frontal cortex; HIPP, hippocampus; MEDU, medulla (specifically inferior olivary nucleus); OCTX, occipital cortex (specifically primary visual cortex); PUTM, putamen; SNIG, substantia nigra; TCTX, temporal cortex; THAL, thalamus; WHMT, intralobular white matter.

### Gene- and pathway-based associations

Gene-level analyses in MAGMA showed a significant enrichment of gene after correcting for multiple testing of the 17854 protein-coding genes. The most significant association was observed for the gene PRDM10 (PR/SET domain 10, 11q24.3) with reading fluency (Z score=4.57, *p*=2.47×10^−6^<2.91×10^−6^=0.05/17184) (supplementary Table S3). In the gene-set analysis of 5497 canonical pathways from KEGG, only a pathway was significantly enriched with reading accuracy in the embryonic germ cell (*p*=4.11×10^−6^<9.10×10^−6^=0.05/5497, β(SE)=0.84(0.19)) (supplementary Table S4). eMAGMA was used to functional tissue-specific eQTL information from GTEx to assign SNPs to genes. In 13 brain regions, we found the most significant association were observed in the cerebellum with VPS33B (location 15q26.1, z score=3.57, *p*=1.74×10^−4^) for reading fluency (supplementary Table S6).

## Discussion

In the present study, we performed GWAS in Chinese children aged 9-11 years-old to identify potential genetic variants associated with reading ability and related cognitive traits. We reported a variant rs6446395 that exceeds genome wide significant association with word reading fluency, which also showed effects on reading accuracy and phonological skills. We identified a gene at the genome-wide level associated with reading fluency in enrichment analyses. Bioinformatics analysis revealed that rs6446395 influenced the expression of EVC in the cerebellar cortex. To our knowledge, it is the first time to report the association of a variant with reading ability at genome-wide significant level in the general population. This variant may be also related to dyslexia.

Reading fluency revealed a genome-wide significant association with a SNP in GWAS and a significant association with a gene (PRDM10) in enrichment analyses, which shows that reading fluency is more genetically affected than other phenotypes. PRDM10 is coding for a transcription factor that contains C2H2-type zinc-fingers which promote nervous system development and involve in the pathogenesis of neuronal storage disease[52].

Firstly, we identified a genome wide significant association effect on word reading fluency. We did not observe any genome wide significant association effect for word reading accuracy, similar to previous GWAS of reading ability[12, 13, 15]. Word reading fluency reflects efficient wording decoding and accounts for a significant proportion of variance in word reading ability[53]. Interestingly, this variant also showed effects on character reading accuracy and phonological skills including RAN, phoneme deletion—behavioral, and cognitive phenotypes important for diagnosis of Chinese developmental dyslexia[28, 54, 55], suggesting its role in dyslexia.

Regarding the biological mechanism of rs6446395, pathway analysis from eMAGMA and eQTL data both suggested that cerebellum may associate with reading fluency, which was consistent with cerebellum theory in dyslexia[56]. Typical readers showed greater activation for faster rates of word presentation in a distributed cortical network including peak activation in cerebellar region. Rs6446395 may affect genes expression in cerebellar to contribute the reading fluency[57]. Mutations of the rs6446395-associated gene EVC are responsible for the EvC (Ellis-van Creveld) syndrome, which is featured with shortened limbs and dental impairments in mutant humans[58, 59]. But the gene expression and protein interaction data suggest that EVC is present in the human central nervous system. However, this is the first study to implicate the genetic variant of EVC in reading fluency. To our knowledge, EVC has not been highlighted in dyslexia studies before, although it was indicated to be related to cognitive impairment.

Regarding the genetics of the cognitive phenotype, although we did not observe genome wide significant effect for any cognitive traits, the most significant association with RAN-color observed in the present study, rs1624971, an intronic SNP located within the CSE1Lgene, are approaching genome wide significance (*p*= 3.48 × 10−7). The assessment of candidate SNPs previously described in dyslexia and related traits also exhibited evidence of replication in the cognitive trait of RAN. Rs2760157 and rs16889506, two SNPs located in the KIAA0319, are associated with RAN-color. Together with the positive results in word reading fluency (speed related task), these results in RAN (speed related task) might again suggested that we should investigate the genetic mechanism of dyslexia in terms of speed related tasks, which may have more vital and more sensitive effects.

Two recent studies in the European populations have identified two variants showing genome wide significant association with RAN[18, 32]. However, none of both SNPs (rs1541518 and rs6963842) withstood correction for multiple testing in our study (RAN dice: *p*=2.52×10-2; RAN picture: *p*=2.00×10-2). The differences between ours and previous studies might be both statistical power and population diversity Alternatively, differences between ours and previous studies might also be caused by task differences. The most significant loci found in previous studies were associated with RAN of letters and RAS of letters and numbers. We could not employ these two tasks in the present study as letters were not familiar items for Chinese elementary school children.

For other weak or no evidence of replication for candidate SNPs, several possible factors may attribute to those inconsistent results. First, different genetic backgrounds of the population analyzed maybe account for the most important factor[60]. Most of those candidate genes and SNPs could be replicated in population speaking English as the native language, but showed nominal or no association with DD in Chinese[61, 62]. Otherwise, the same SNPs in different populations could have contrasting direction of effects. Second, the effects of candidate SNPs might be lower in the current normal population due to small variance, while these effects might be replicated in Chinese DD samples, which might be valuable for further investigation. Also, environmental factors, in particular preschool education and social economic status were different between areas and might lead to variations between studies[63].

For the limitations of the study, sample size is an obvious one compared with genome wide studies for chronic diseases such as diabetes or schizophrenia[64]. In addition, this study was not replicated in other cohorts, limiting the identification of biological clues surrounding the aetiology of a complex disorder. Obviously, our brain eQTL analysis used the UK database, and the expression of EVC may be biased due to cohort differences between Asian and UK database. It should also be noted that eMAGMA and eQTL data do not align on the same SNP to gene pairing, which might require further investiation in future studies. Despite limitations above, the strength of the current research is for the first time perform a GWAS of the reading and related cognitive traits in Chinese children genotyped by using Illumina Asian Screening Array, which is specially designed for East Asian population.

In conclusion, we identified rs6446395 in EVC at the region of chromosome 4p16.2 that showed genome wide significant association effect with word reading fluency in a sample of Chinese children. This is the first evidence that EVC has an impact on word reading fluency. Overall, this study contributed to broaden the genetic basis of reading ability in Chinese children and developmental dyslexia.

## Acknowledgments

This work was funded by National Natural Science Foundation of China (61807023), Funds for Humanities and Social Sciences Research of the Ministry of Education (CN) (17XJC190010), Natural Science Foundation of Shaanxi Province (CN) (2018JQ8015), and Fundamental Research Funds for the Central Universities (CN) (GK201702011) to Jingjing Zhao. This study was also funded by Fund for Humanities and Social Sciences Research of the Ministry of Education (19YJC190023), and the China Postdoctoral Science Foundation funding project (2019M663924XB), Natural Science Foundation of Shaanxi Province (CN) (2021JQ-309), and Planning Subject for the 14th Five Year Plan of Shaanxi Education Sciences (SGH21Y0040) to Zhengjun Wang.

## Conflict of interest

The authors have no conflicts of interest to disclose.

## Figure legends

**Figure S1.**
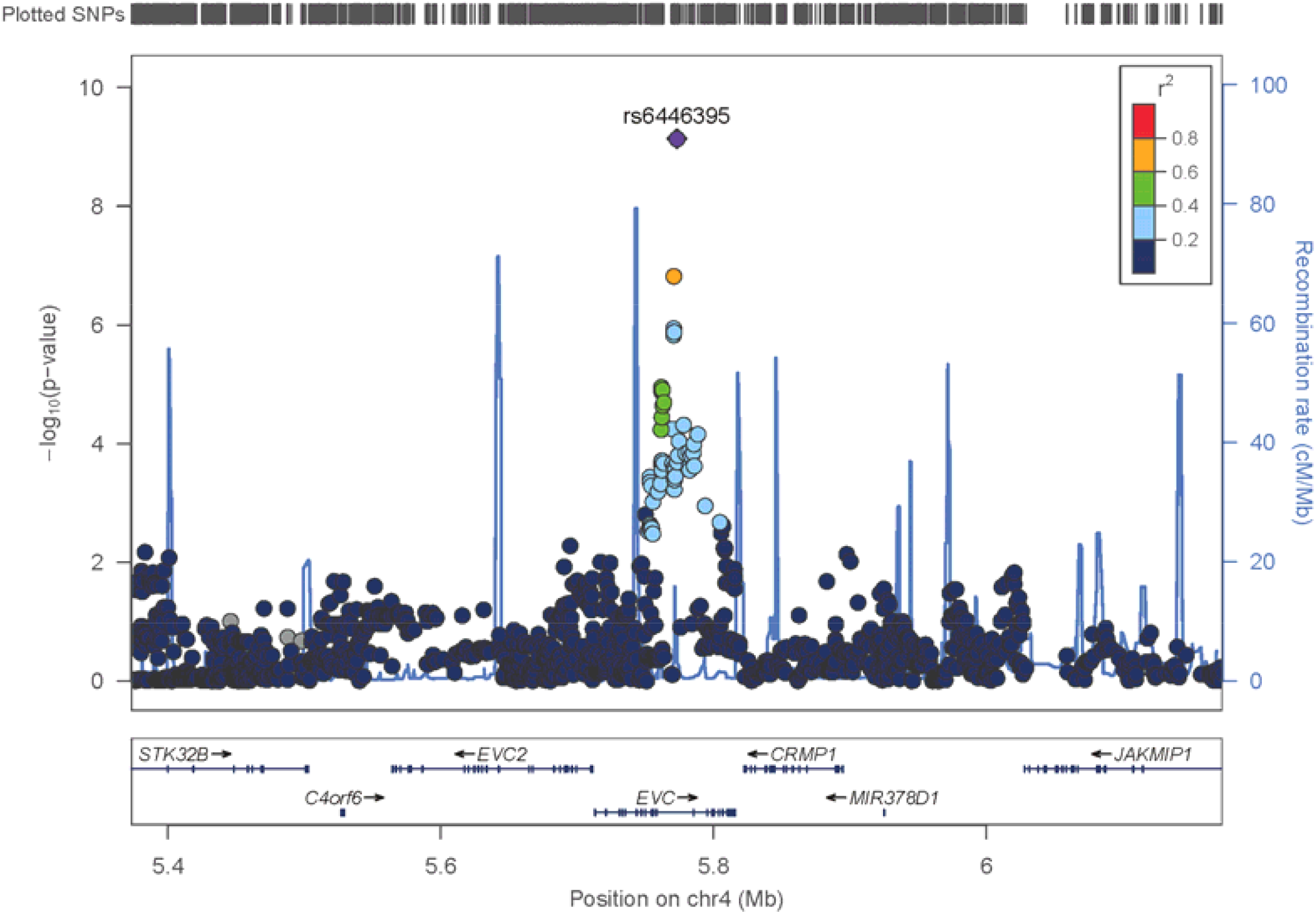
Regional plot for the significant locus rs6446395 for reading fluency. The most significantly associated variants are highlighted in violet. Plots were made using LocusZoom v0.4.8

**Figure S2.**
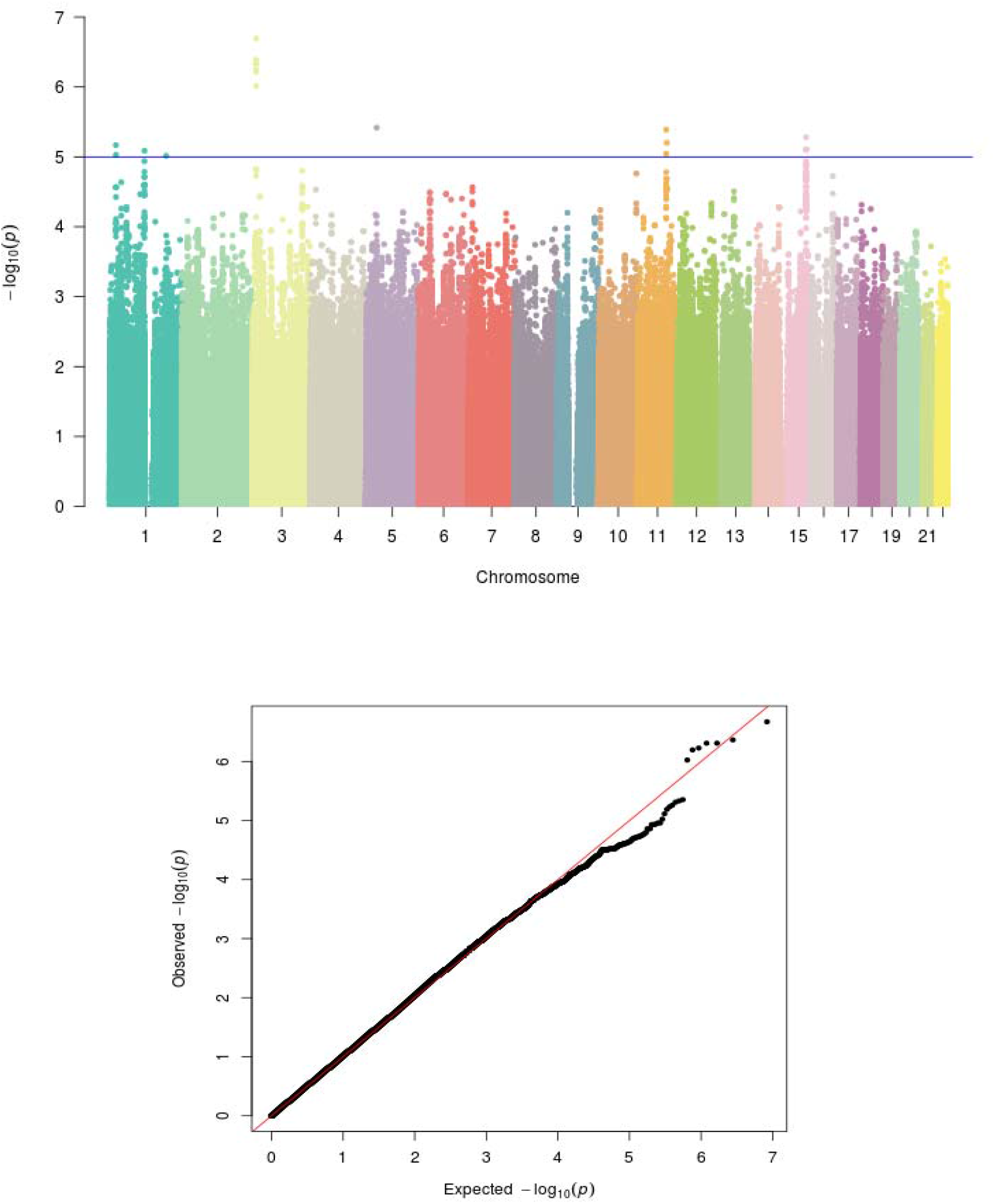
Manhattan plot summarising the results and QQ plots of the genome-wide association study (GWAS) for reading accuracy. The blue line represent the genome wide significance threshold (*p*<1×10^−5^). λ= 1.021

**Figure S3.**
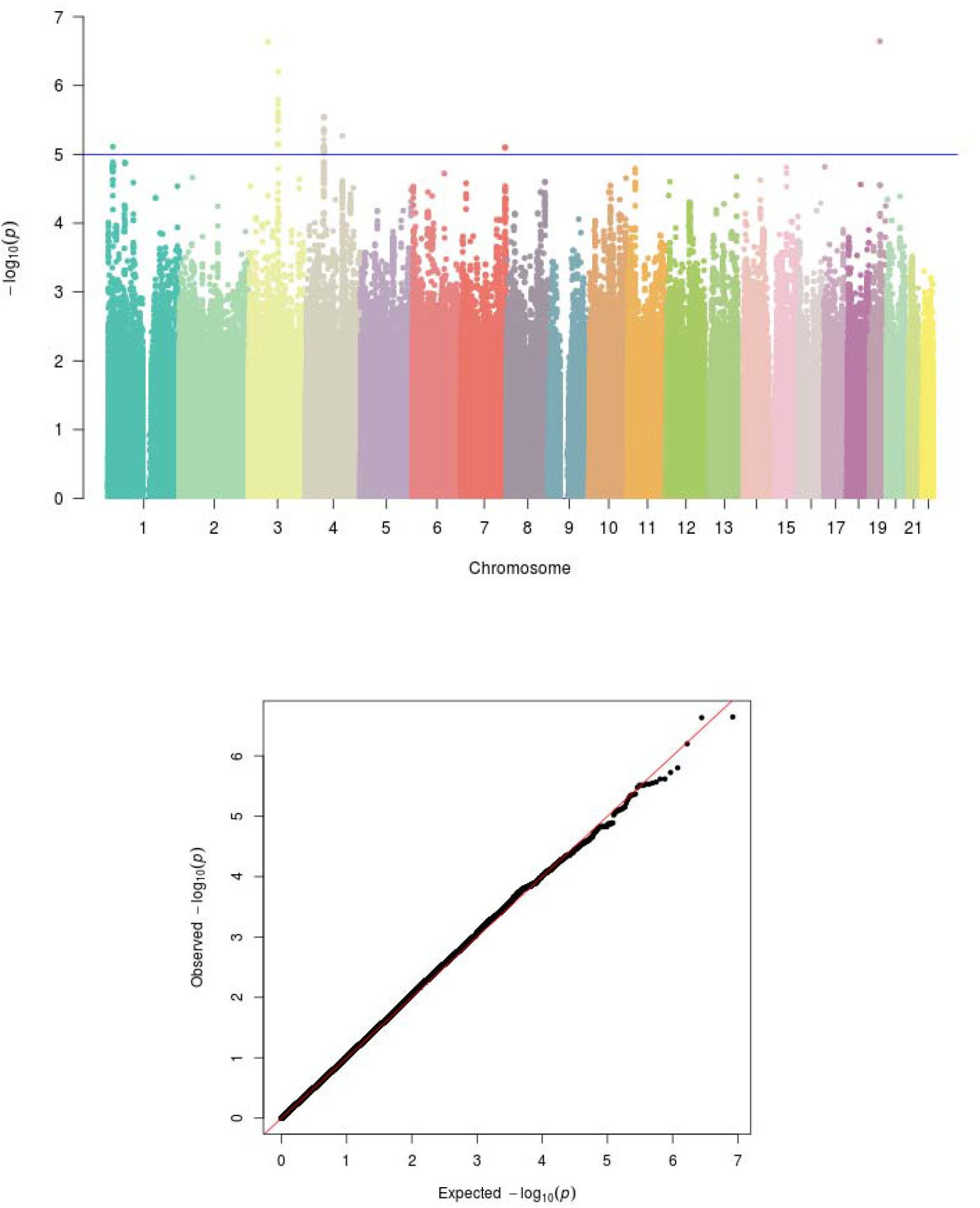
Manhattan plot summarising the results and QQ plots of the genome-wide association study (GWAS) for Phoneme awareness. The blue line represent the genome wide significance threshold (*p*<1×10^−5^). λ= 1.013

**Figure S4.**
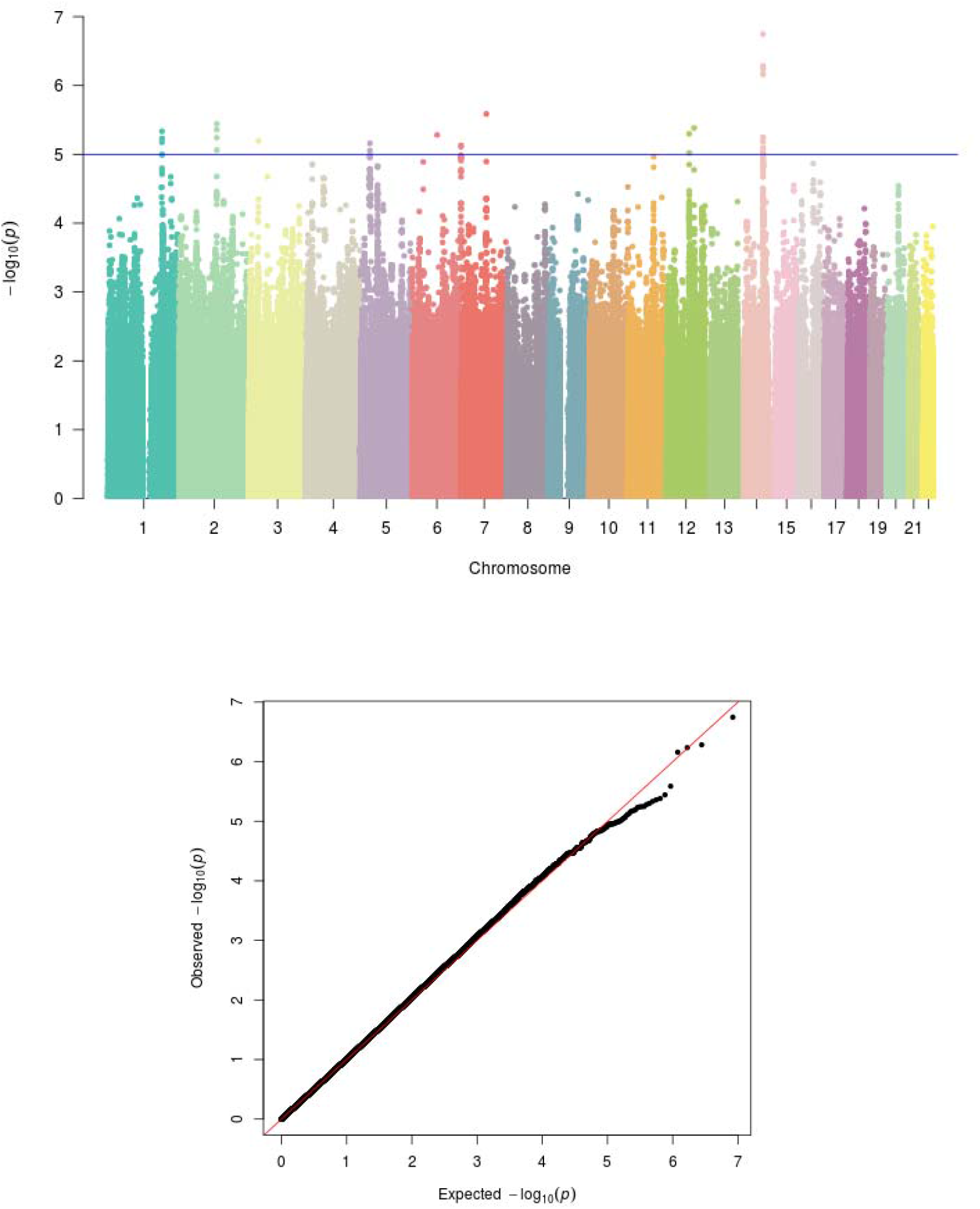
Manhattan plot summarising the results and QQ plots of the genome-wide association study (GWAS) for Morphological awareness. The blue line represent the genome wide significance threshold (*p*<1×10^−5^). λ= 1.003

**Figure S5.**
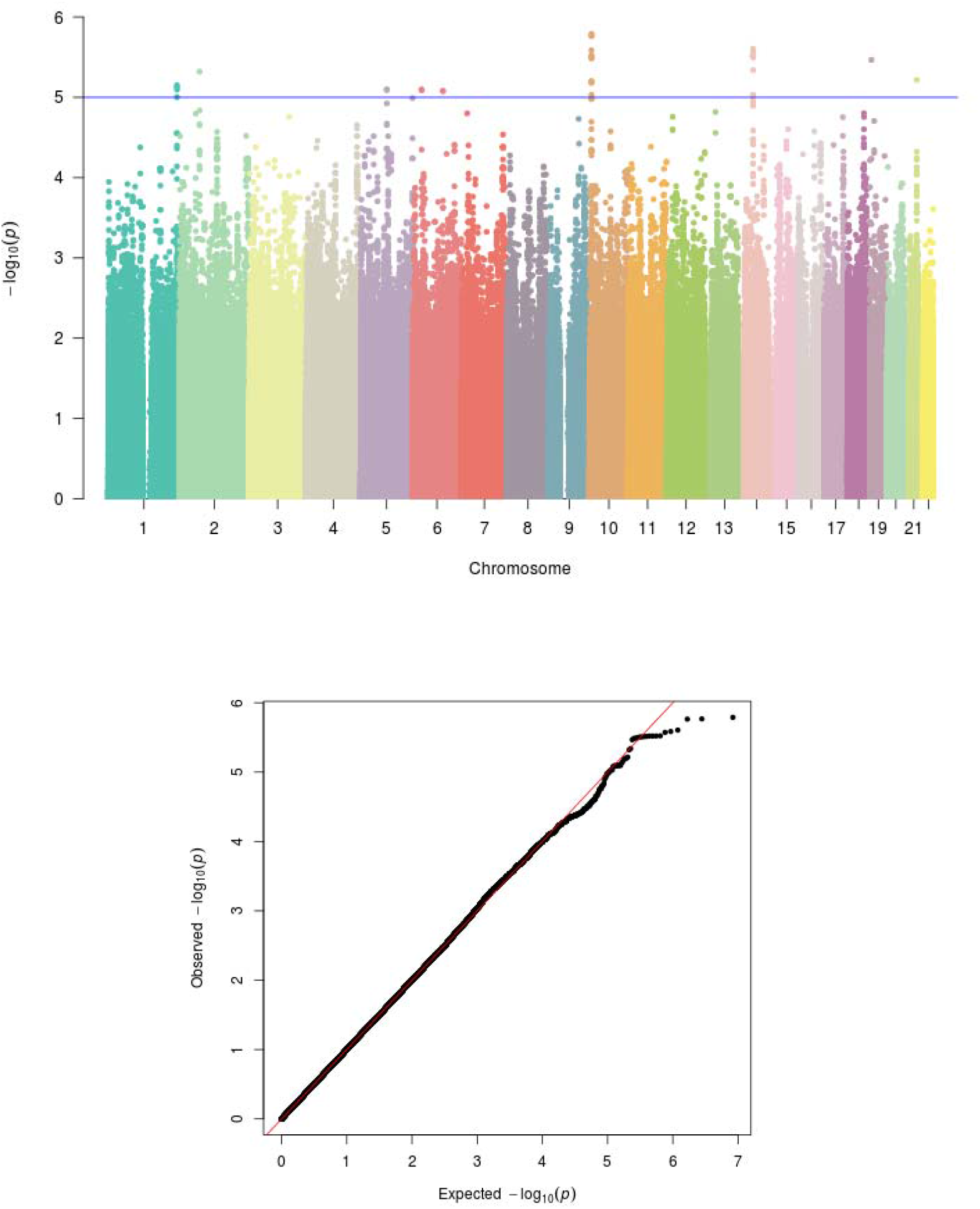
Manhattan plot summarising the results and QQ plots of the genome-wide association study (GWAS) for RAN-digit. The blue line represent the genome wide significance threshold (*p*<1×10^−5^). λ= 1.004

**Figure S6.**
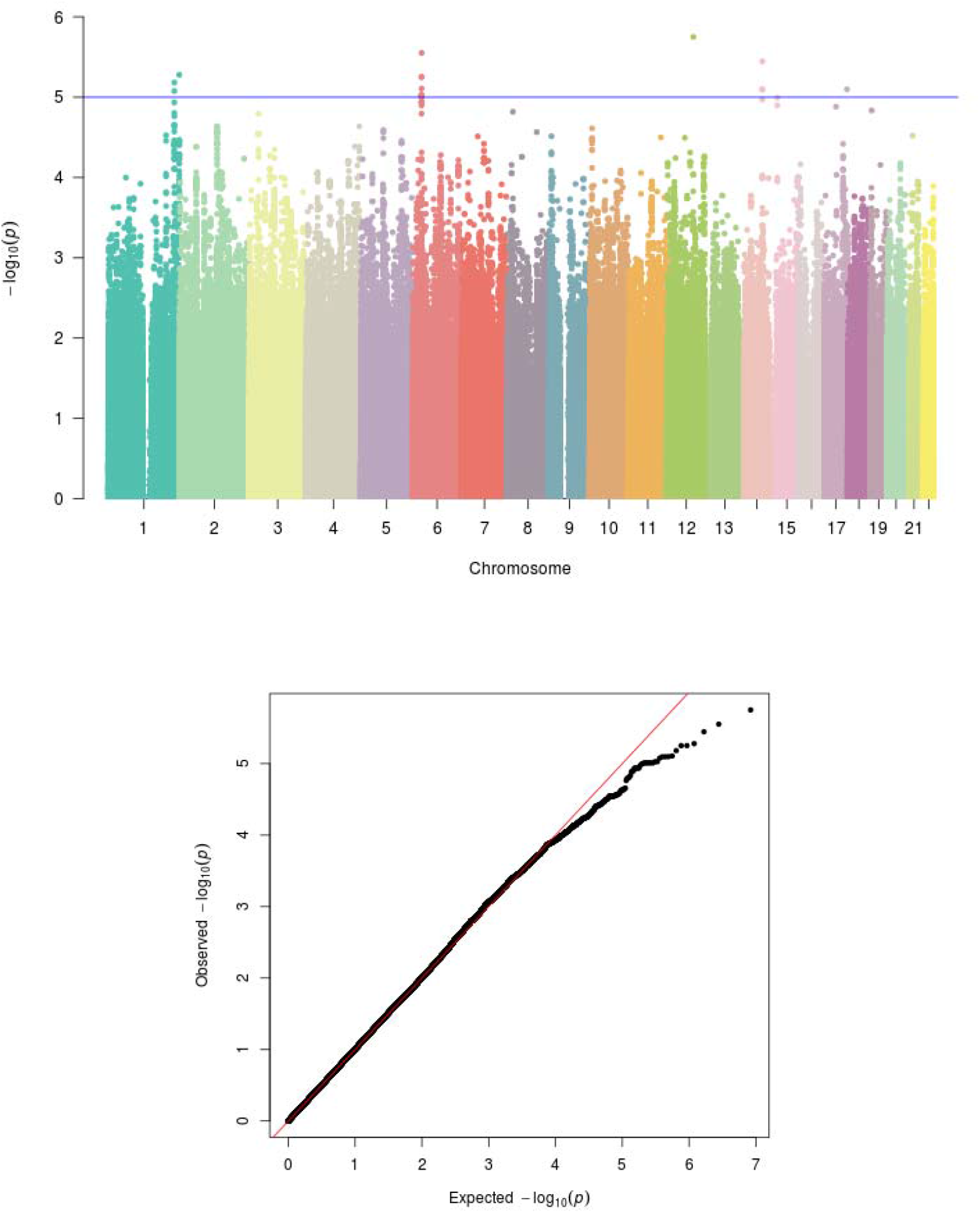
Manhattan plot summarising the results and QQ plots of the genome-wide association study (GWAS) for RAN-dic. The blue line represent the genome wide significance threshold (*p*<1×10^−5^). λ= 1.01

**Figure S7.**
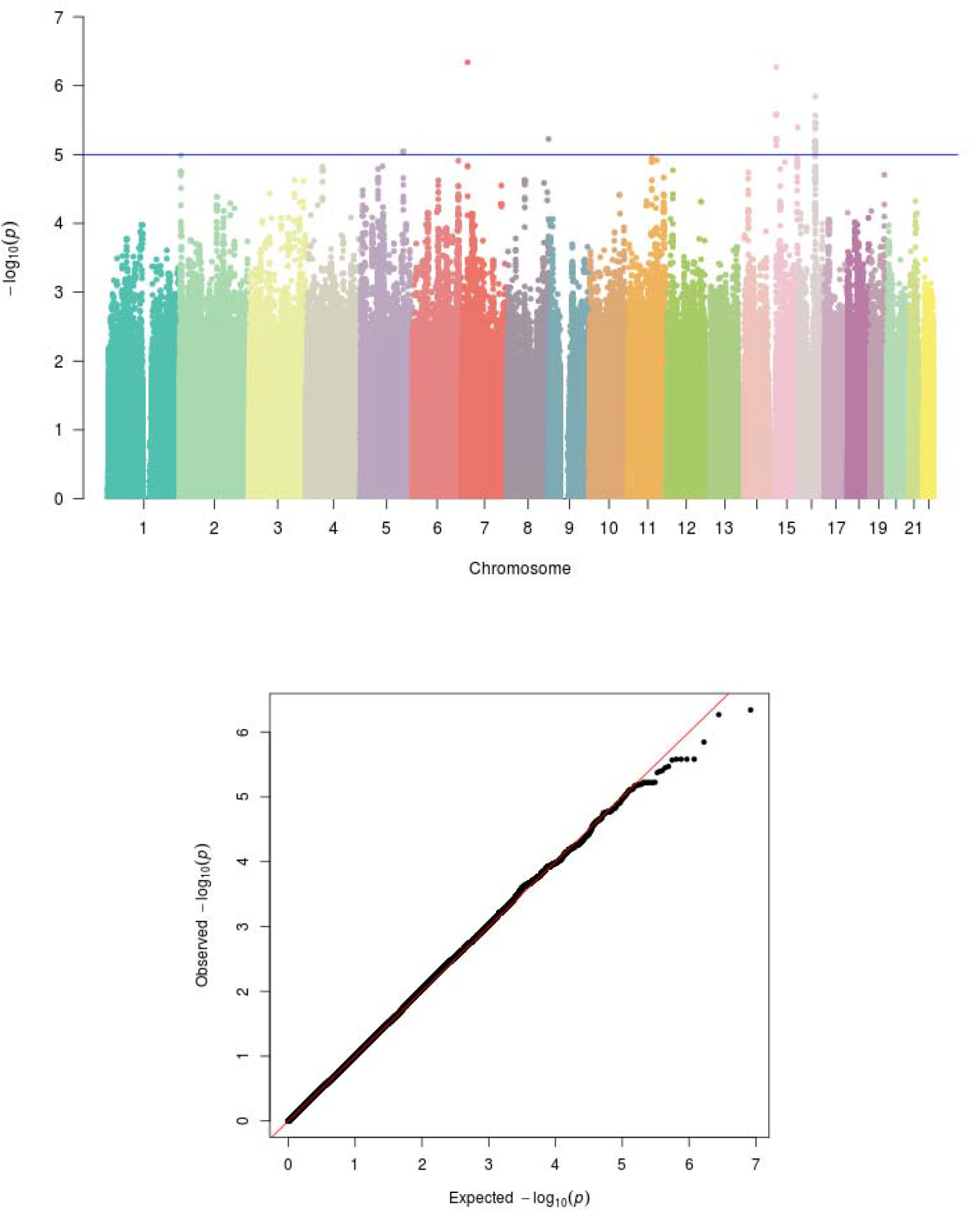
Manhattan plot summarising the results and QQ plots of the genome-wide association study (GWAS) for RAN-picture. The blue line represent the genome wide significance threshold (*p*<1×10^−5^). λ= 1.011

**Figure S8.**
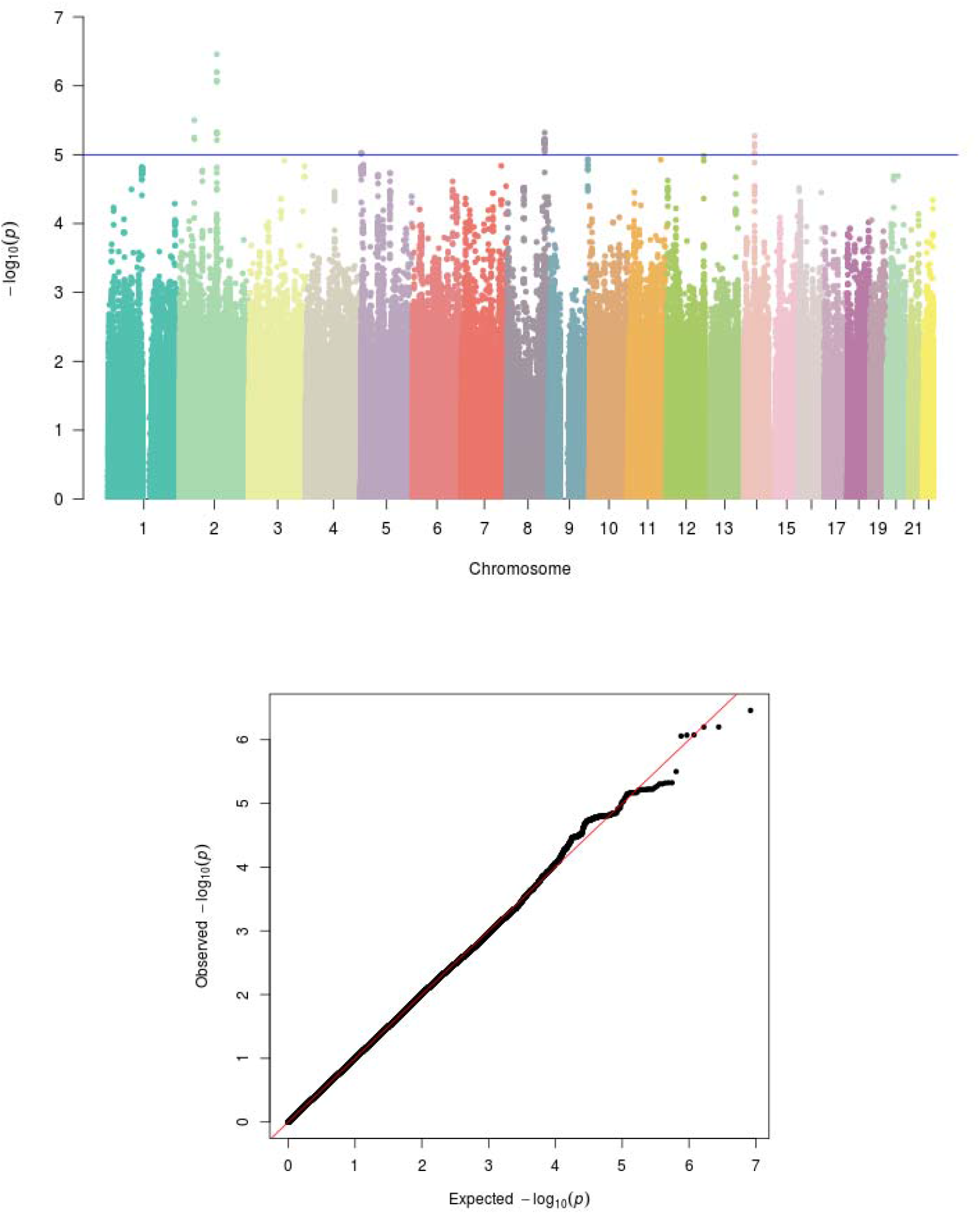
Manhattan plot summarising the results and QQ plots of the genome-wide association study (GWAS) for RAN-color. The blue line represent the genome wide significance threshold (*p*<1×10^−5^). λ= 1.009

**Figure S9.**
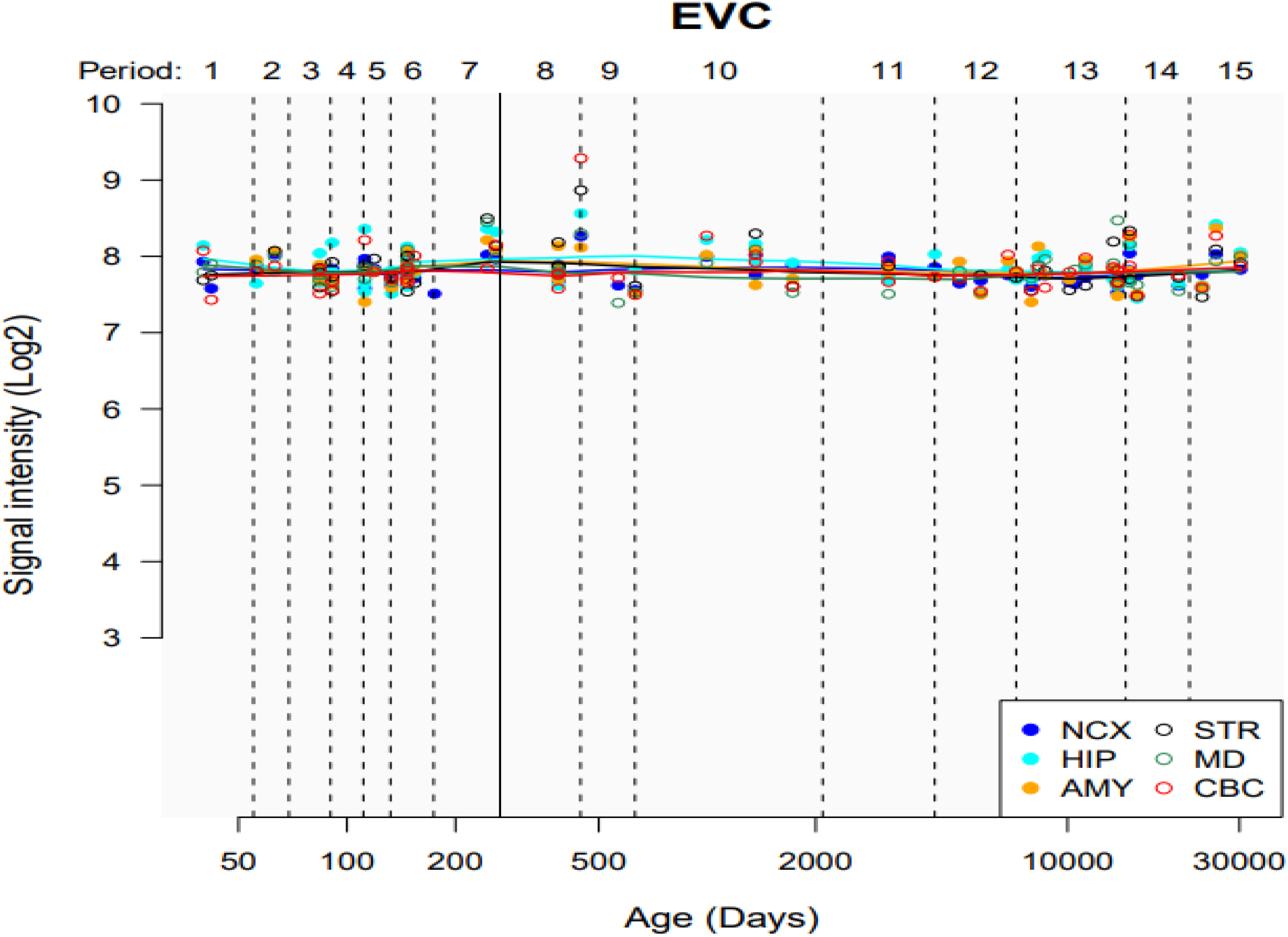
Temporal and regional human EVC expression

**Figure S10.**
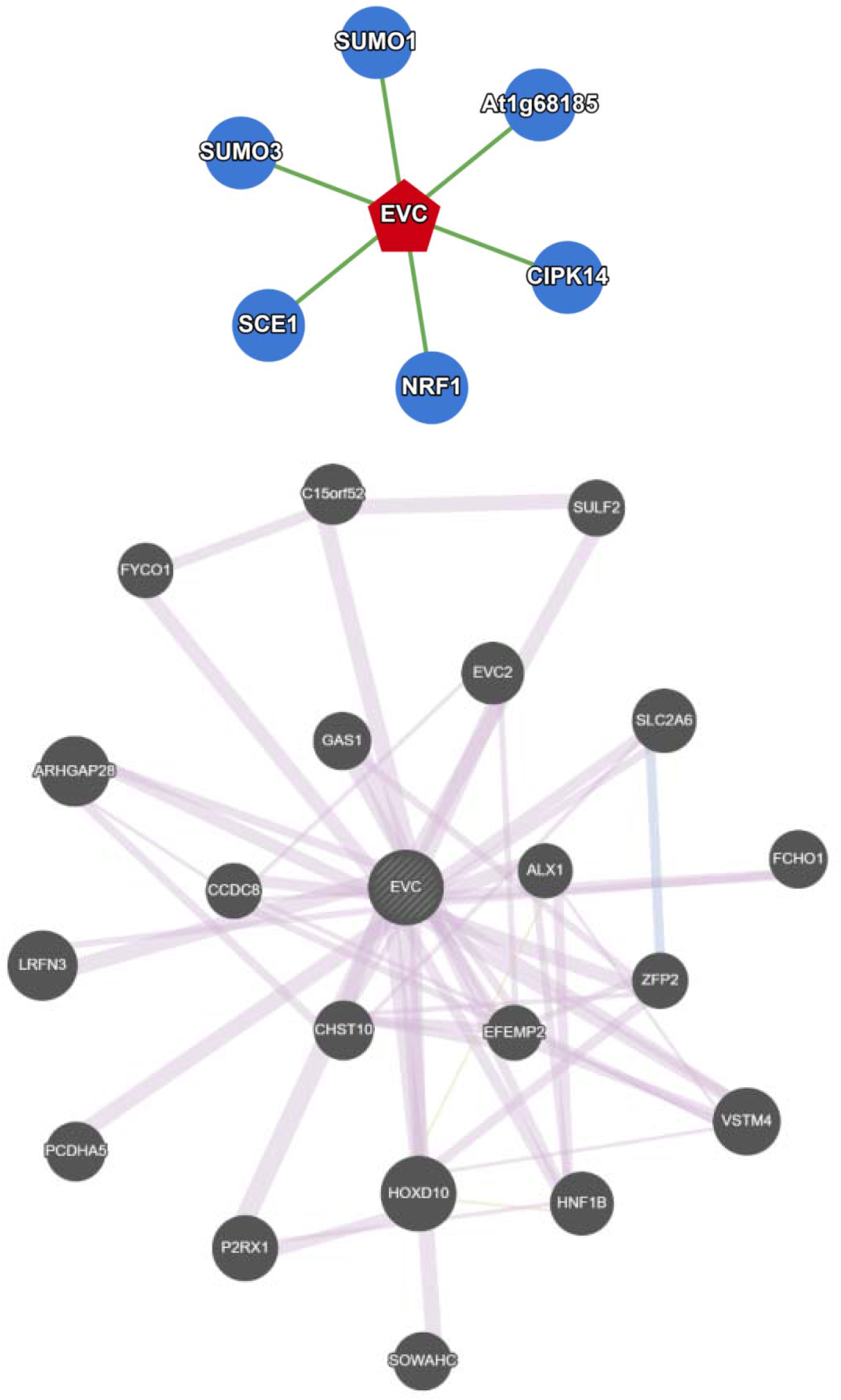
Six interacting genes and 20 co-expressed genes were observed indicating their association with EVC.

## Notes

### Competing Interest Statement

The authors have declared no competing interest.

