## Supplemental Methods for "A genome wide association study identifies a new variant associated with word reading fluency in Chinese children"

**Supplementary Methods**

***Word reading accuracy***

Chinese character recognition test was employed to measure each child’s word reading accuracy ^1-3^. The test consisted of 150 single Chinese characters selected from China’s *Elementary School Textbooks* (1996). The average frequency of the characters was 182 per million (ranging from 0 to 2282), and the reliability of this test was 0.95 ^2^. Each child was individually tested and required to read aloud each character at a time.

***Word reading fluency***

Word list reading task ^2^ was used to measure each child’s word reading fluency. In this task, children were asked to name a list of 180 two character words as rapidly and accurately as possible. All these words were from primary school text books and have been learned before grade 3, such as “我们(we)” and “太阳(sun)”. The mean frequency of these words was 212.77 per million ^4^. Since words included in this task were all simple, this task was administrated to test children’s word reading fluency. The total time for naming the whole word list was recorded as measurement of word reading fluency.

***Phoneme awareness***

In this task, the experimenter first orally presented a one-syllable real word to each child. The child was asked to take away a given phoneme from the syllable and speak out the rest of the syllable. The task included 16 items, with 2 initial phoneme deletion items (e.g., /mei4/(sister) without the /m/), 4 middle phoneme deletion items (e.g., /tuan2/(group) without the /u/), and 10 final phoneme deletion items (e.g., /guan1/(close) without the /n/). This task has been widely used in previous Chinese reading development and impairment studies ^5-7^. The reliability (Chronbach’s alpha) of the test in the present study was 0.90.

***Rapid automatized naming***

Rapid automatized naming includes four tasks: *rapid automatized digit naming* ^5^, *rapid automatized picture naming* ^8^, *rapid automatized dice naming* ^9^, and *rapid automatized color naming* ^8^. Four series of 40 items (digits, pictures, dices, and colors) were presented to each child, with each type of items on a separate sheet of paper. The digits (2, 4, 6, 7, and 9) were used as stimuli of rapid automatized digit naming task. The pictures (dog, flower, book, shoe, and window) were used as stimuli of rapid automatized picture naming task. Pictures of dices (one, two, three, four, and five) were used as stimuli of rapid automatized dice naming task. The colors (red, yellow, black, green, and blue) were used as stimuli of rapid automatized color naming task. Each sheet includes eight rows with five items in a row. Children were required to name each type of items as rapidly as possible. Each child named twice for each sheet. The measurement of rapid automatized naming was the average naming times for the two times of each type of items. The test-retest reliabilities of rapid automatized digit, picture, dice, and color naming tasks were 0.87, 0.82, 0.74, and 0.74, respectively.

***Morphological awareness***

In this task, administered individually to each child, the experimenter orally presented a two-syllable Chinese word. Within the two-morpheme word, one morpheme was identified ^5^. The child was then asked to produce two words with the target morpheme. One of the morphemes was supposed to have the same meaning as the target morpheme. The other morpheme was supposed to have a meaning different from its original meaning. However, both morphemes were identical in pronunciation. For example, when the experimenter gave the word bei1bao1 (meaning backpack), the child was asked to produce a new word with the morpheme [bao1] in which the [bao1] had the same meaning as it did in bei1bao1. One acceptable answer would be shu1bao1 (meaning school bag). The child was also asked to say a word that included the morpheme [bao1] in which its meaning was different from that in bei1bao1. An example is bao1zi1 (meaning bun). All items consisted of real words. There were 30 items in total.
