## Supplemental Table 1 for "A genome wide association study identifies a new variant associated with word reading fluency in Chinese children"

Table S1. Pearson correlation coefficients across character recognition, reading fluency, phoneme awareness, morphological awareness, and RAN dig/dic/pict/color.

|  | RC | RF | PD | MA | RAN-digit | RAN-dice | RAN-picture | RAN-color |
| --- | --- | --- | --- | --- | --- | --- | --- | --- |
| RC | 1 |  |  |  |  |  |  |  |
| RF | -0.562 | 1 |  |  |  |  |  |  |
| PD | 0.379 | -.324 | 1 |  |  |  |  |  |
| MA | 0.373 | -.318 | .397 | 1 |  |  |  |  |
| RAN-digit | -.435 | .611 | -.272 | -.239 | 1 |  |  |  |
| RAN-dice | -.396 | .461 | -.256 | -.218 | .595 | 1 |  |  |
| RAN-picture | -.398 | .468 | -.241 | -.241 | .512 | .615 | 1 |  |
| RAN-color | -.339 | .422 | -.252 | -.223 | .496 | .584 | .661 | 1 |

Note: All *p*<1e^-26^ (two-tailed); RC: reading character; RF: reading fluency; PD: phoneme deletion (phoneme awareness); MA: morphological awareness.
