## Supplemental Table 2 for "A genome wide association study identifies a new variant associated with word reading fluency in Chinese children"

| LeadingSNPs | SNP | R^2^ | MAF | T/I | GENE | Identified by | RFluency_BETA | P |
| --- | --- | --- | --- | --- | --- | --- | --- | --- |
| rs3756821 |  |  | 0.288 | G | KIAA0319 | Zhao et al^[45]^ | 1.356 | 0.1918 |
|  | rs3212236 | .493 | 0.451 | I | KIAA0319 | Harold et al^[65]^ | 1.411 | 0.132 |
|  | rs16889556 | .454 | 0.157 | I | KIAA0319 | Zhao et al^[45]^ | 1.135 | 0.3841 |
|  | rs6935076 | .721 | 0.228 | G | KIAA0319 | Cope et al^[66]^ | 1.82 | 0.1074 |
| rs761100 |  |  | 0.171 | G | KIAA0319 | Harold et al^[65]^ | 0.644 | 0.6055 |
|  | rs2038137 | 0.927 | 0.161 | I | KIAA0319 | Zhao et al^[45]^ | 0.7229 | 0.5735 |
|  | rs4504469 | 0.626 | 0.175 | G | KIAA0319 | Francks et al^[67]^ | 0.7664 | 0.5308 |
|  | rs2179515 | 0.914 | 0.16 | I | KIAA0319 | Harold et al^[65]^ | -0.3273 | 0.8123 |
| rs807507 |  |  | 0.246 | I | KIAA0319 | Zhao et al^[45]^ | -0.06696 | 0.9504 |
|  | rs12193738 | 0.971 | 0.246 | I | KIAA0319 | Zhao et al^[45]^ | -0.7152 | 0.5056 |
|  | rs3903801 | 0.858 | 0.257 | I | KIAA0319 | Zhao et al^[45]^ | -1.041 | 0.3232 |
| rs1000585 |  |  | 0.422 | I | MRPL19 | Anthoni et al^[68]^ | 0.5807 | 0.5402 |
|  | rs917235 | 0.944 | 0.424 | I | MRPL19 | Anthoni et al^[68]^ | 0.3508 | 0.5402 |
| rs3743204 |  |  | 0.16 | I | DYX1C1 | Wigg et al^[69]^ | -1.182 | 0.3654 |
|  | rs600753 | 0.542 | 0.246 | I | DYX1C1 | Scerri et al^[70]^ | -0.7959 | 0.4739 |
| rs807724 |  | 1.00 | 0.056 | G | DCDC2 | Schumacher et al^[71]^ | -0.247 | 0.9049 |
| rs807701 |  | 1.00 | 0.246 | G | DCDC2 | Schumacher et al^[71]^ | -1.297 | 0.2382 |
| rs2143340 |  | 1.00 | 0.171 | G | KIAA0319 | Francks et al^[67]^ | 0.009202 | 0.9421 |
| rs16889506 |  | 1.00 | 0.167 | I | KIAA0319 | Zhao et al^[45]^ | -1.779 | 0.1633 |
| rs699463 |  | 1.00 | 0.14 | G | KIAA0319 | Zhao et al^[45]^ | -0.3273 | 0.8123 |
| rs9366577 |  | 1.00 | 0.052 | G | KIAA0319 | Zhao et al^[45]^ | 2.574 | 0.2315 |
| rs793862 |  | 1.00 | 0.389 | G | DCDC2 | Schumacher et al^[71]^ | 0.01535 | 0.9874 |
| rs714939 |  | 1.00 | 0.398 | I | MRPL19 | Anthoni et al^[68]^ | -0.7081 | 0.4626 |
| rs16889556 |  | 1.00 | 0.213 | G | KIAA0319 | Zhao et al^[45]^ | 0.9126 | 0.4352 |
| rs6732511 |  | 1.00 | 0.072 | G | MRPL19 | Anthoni et al^[68]^ | -0.1563 | 0.9312 |
| rs2760157 |  | 1.00 | 0.438 | G | KIAA0319 | Zhao et al^[45]^ | 0.5783 | 0.5352 |
| rs2038139 |  |  | 0.02 | I | KIAA0319 | Zhao et al^[45]^ |  |  |
| rs6963842 |  | 1.00 | 0.209 | G | LAMB1 | Truong et al^[32]^ | -0.4032 | 0.7299 |
| rs11177505 |  |  |  | No present | Null | Truong et al^[32]^ |  |  |
| rs4320486 |  |  |  | No present | Null | Truong et al^[32]^ |  |  |
| rs17663182 |  |  |  | No present | Null | Gialluisi et al^[18]^ |  |  |
| rs17605546 |  |  |  | No present | Null | Gialluisi et al^[18]^ |  |  |
| rs34822091 |  |  |  | No present | Null | Gialluisi et al^[18]^ |  |  |
| rs16928927 |  |  |  | No present | Null | Gialluisi et al^[18]^ |  |  |
| rs4571421 |  | 1.00 | 0.427 | I | LINC02118 | Gialluisi et al^[18]^ | 1.02 | 0.2851 |
| rs76161559 |  |  |  | No present | Null | Gialluisi et al^[18]^ |  |  |
| rs4307051 |  |  |  | No present | Null | Gialluisi et al^[18]^ |  |  |
| rs200580547 |  |  |  | No present | Null | Gialluisi et al^[18]^ |  |  |

| LeadingSNPs | SNP | R^2^ | MAF | GENE | Identified by | Raccuracy_BETA | P |
| --- | --- | --- | --- | --- | --- | --- | --- |
| rs3756821 |  |  | 0.288 | KIAA0319 | Zhao et al^[32]^ | 0.6031 | 0.3235 |
|  | rs3212236 | .493 | 0.451 | KIAA0319 | Harold et al^[52]^ | -0.2771 | 0.6148 |
|  | rs16889556 | .454 | 0.157 | KIAA0319 | Zhao et al^[32]^ | 0.7115 | 0.3538 |
|  | rs6935076 | .721 | 0.228 | KIAA0319 | Cope et al^[53]^ | 0.6094 | 0.3603 |
| rs761100 |  |  | 0.171 | KIAA0319 | Harold et al^[52]^ | -1.641 | 0.02501 |
|  | rs2038137 | 0.927 | 0.161 | KIAA0319 | Zhao et al^[32]^ | -1.468 | 0.05177 |
|  | rs4504469 | 0.626 | 0.175 | KIAA0319 | Francks et al.^[54]^ | -1.434 | 0.04574 |
|  | rs2179515 | 0.914 | 0.16 | KIAA0319 | Harold et al^[52]^ | -0.0617 | 0.9392 |
| rs807507 |  |  | 0.246 | KIAA0319 | Zhao et al^[32]^ | -0.3506 | 0.5794 |
|  | rs12193738 | 0.971 | 0.246 | KIAA0319 | Zhao et al^[32]^ | -0.1033 | 0.8702 |
|  | rs3903801 | 0.858 | 0.257 | KIAA0319 | Zhao et al^[32]^ | -0.01301 | 0.9833 |
| rs1000585 |  |  | 0.422 | MRPL19 | Anthoni et al.^[55]^ | 0.2294 | 0.6801 |
|  | rs917235 | 0.944 | 0.424 | MRPL19 | Anthoni et al.^[55]^ | 0.3666 | 0.5119 |
| rs3743204 |  |  | 0.16 | DYX1C1 | Wigg et al.^[56]^ | -0.4823 | 0.5291 |
|  | rs600753 | 0.542 | 0.246 | DYX1C1 | Scerri et al.^[58]^ | -0.7784 | 0.2331 |
| rs807724 |  | 1.00 | 0.056 | DCDC2 | Schumacher et al^[57]^ | 0.5014 | 0.6797 |
| rs807701 |  | 1.00 | 0.246 | DCDC2 | Schumacher et al^[57]^ | 0.3816 | 0.5547 |
| rs2143340 |  | 1.00 | 0.171 | KIAA0319 | Francks et al.^[54]^ | 0.3078 | 0.6799 |
| rs16889506 |  | 1.00 | 0.167 | KIAA0319 | Zhao et al^[32]^ | 1.115 | 0.1373 |
| rs699463 |  | 1.00 | 0.14 | KIAA0319 | Zhao et al^[32]^ | -0.0617 | 0.9392 |
| rs9366577 |  | 1.00 | 0.052 | KIAA0319 | Zhao et al^[32]^ | 0.3192 | 0.8013 |
| rs793862 |  | 1.00 | 0.389 | DCDC2 | Schumacher et al^[57]^ | -0.9319 | 0.1019 |
| rs714939 |  | 1.00 | 0.398 | MRPL19 | Anthoni et al.^[55]^ | 0.8222 | 0.1465 |
| rs1091031 |  | 1.00 | 0.213 | KIAA0319 | Zhao et al^[32]^ | -0.09686 | 0.8897 |
| rs6732511 |  | 1.00 | 0.072 | MRPL19 | Anthoni et al.^[55]^ | 0.9021 | 0.396 |
| rs2760157 |  | 1.00 | 0.438 | KIAA0319 | Zhao et al^[32]^ | -0.003452 | 0.995 |
| rs6963842 |  | 1.00 | 0.209 | LAMB1 | Truong et al^[21]^ | 0.0308 | 0.9642 |
| rs4571421 |  | 1.00 | 0.427 | LINC02118 | Gialluisi et al | -0.4803 | 0.392 |

| LeadingSNPs | SNP | R^2^ | MAF | GENE | Identified by | PA_BETA | P |
| --- | --- | --- | --- | --- | --- | --- | --- |
| rs3756821 |  |  | 0.288 | KIAA0319 | Zhao et al^[32]^ | 0.1021 | 0.3124 |
|  | rs3212236 | .493 | 0.451 | KIAA0319 | Harold et al^[52]^ | 0.07447 | 0.4137 |
|  | rs16889556 | .454 | 0.157 | KIAA0319 | Zhao et al^[32]^ | 0.04293 | 0.7353 |
|  | rs6935076 | .721 | 0.228 | KIAA0319 | Cope et al^[53]^ | 0.04467 | 0.685 |
| rs761100 |  |  | 0.171 | KIAA0319 | Harold et al^[52]^ | -0.01471 | 0.9036 |
|  | rs2038137 | 0.927 | 0.161 | KIAA0319 | Zhao et al^[32]^ | -0.02558 | 0.8381 |
|  | rs4504469 | 0.626 | 0.175 | KIAA0319 | Francks et al.^[54]^ | -0.1341 | 0.26 |
|  | rs2179515 | 0.914 | 0.16 | KIAA0319 | Harold et al^[52]^ | -0.04156 | 0.7394 |
| rs807507 |  |  | 0.246 | KIAA0319 | Zhao et al^[32]^ | 0.02883 | 0.7827 |
|  | rs12193738 | 0.971 | 0.246 | KIAA0319 | Zhao et al^[32]^ | 0.04165 | 0.69 |
|  | rs3903801 | 0.858 | 0.257 | KIAA0319 | Zhao et al^[32]^ | 0.0385 | 0.707 |
| rs1000585 |  |  | 0.422 | MRPL19 | Anthoni et al.^[55]^ | 0.001165 | 0.9899 |
|  | rs917235 | 0.944 | 0.424 | MRPL19 | Anthoni et al.^[55]^ | 0.02459 | 0.7904 |
| rs3743204 |  |  | 0.16 | DYX1C1 | Wigg et al.^[56]^ | -0.03232 | 0.7996 |
|  | rs600753 | 0.542 | 0.246 | DYX1C1 | Scerri et al.^[58]^ | -0.003859 | 0.9716 |
| rs807724 |  | 1.00 | 0.056 | DCDC2 | Schumacher et al^[57]^ | -0.1633 | 0.4158 |
| rs807701 |  | 1.00 | 0.246 | DCDC2 | Schumacher et al^[57]^ | 0.01269 | 0.9056 |
| rs2143340 |  | 1.00 | 0.171 | KIAA0319 | Francks et al.^[54]^ | -0.1039 | 0.4001 |
| rs16889506 |  | 1.00 | 0.167 | KIAA0319 | Zhao et al^[32]^ | 0.1714 | 0.168 |
| rs699463 |  | 1.00 | 0.14 | KIAA0319 | Zhao et al^[32]^ | -0.1107 | 0.409 |
| rs9366577 |  | 1.00 | 0.052 | KIAA0319 | Zhao et al^[32]^ | 0.01088 | 0.9588 |
| rs793862 |  | 1.00 | 0.389 | DCDC2 | Schumacher et al^[57]^ | -0.1452 | 0.124 |
| rs714939 |  | 1.00 | 0.398 | MRPL19 | Anthoni et al.^[55]^ | -0.001463 | 0.9876 |
| rs1091031 |  | 1.00 | 0.213 | KIAA0319 | Zhao et al^[32]^ | -0.05949 | 0.6003 |
| rs6732511 |  | 1.00 | 0.072 | MRPL19 | Anthoni et al.^[55]^ | -0.1097 | 0.5334 |
| rs2760157 |  | 1.00 | 0.438 | KIAA0319 | Zhao et al^[32]^ | -0.00804 | 0.9295 |
| rs6963842 |  | 1.00 | 0.209 | LAMB1 | Truong et al^[21]^ | 0.04987 | 0.6616 |
| rs4571421 |  | 1.00 | 0.427 | LINC02118 | Gialluisi et al | 0.01626 | 0.8614 |

| LeadingSNPs | SNP | R^2^ | MAF | GENE | Identified by | MA_BETA | P |
| --- | --- | --- | --- | --- | --- | --- | --- |
| rs3756821 |  |  | 0.288 | KIAA0319 | Zhao et al^[32]^ | -0.03867 | 0.8286 |
|  | rs3212236 | .493 | 0.451 | KIAA0319 | Harold et al^[52]^ | -0.007869 | 0.9611 |
|  | rs16889556 | .454 | 0.157 | KIAA0319 | Zhao et al^[32]^ | 0.1507 | 0.5015 |
|  | rs6935076 | .721 | 0.228 | KIAA0319 | Cope et al^[53]^ | -0.07152 | 0.713 |
| rs761100 |  |  | 0.171 | KIAA0319 | Harold et al^[52]^ | 0.1011 | 0.641 |
|  | rs2038137 | 0.927 | 0.161 | KIAA0319 | Zhao et al^[32]^ | 0.01997 | 0.9288 |
|  | rs4504469 | 0.626 | 0.175 | KIAA0319 | Francks et al.^[54]^ | -0.2133 | 0.3147 |
|  | rs2179515 | 0.914 | 0.16 | KIAA0319 | Harold et al^[52]^ | -0.05139 | 0.8178 |
| rs807507 |  |  | 0.246 | KIAA0319 | Zhao et al^[32]^ | 0.2693 | 0.1476 |
|  | rs12193738 | 0.971 | 0.246 | KIAA0319 | Zhao et al^[32]^ | 0.2989 | 0.1072 |
|  | rs3903801 | 0.858 | 0.257 | KIAA0319 | Zhao et al^[32]^ | 0.223 | 0.2205 |
| rs1000585 |  |  | 0.422 | MRPL19 | Anthoni et al.^[55]^ | 0.06588 | 0.6853 |
|  | rs917235 | 0.944 | 0.424 | MRPL19 | Anthoni et al.^[55]^ | 0.08274 | 0.6125 |
| rs3743204 |  |  | 0.16 | DYX1C1 | Wigg et al.^[56]^ | 0.08075 | 0.7192 |
|  | rs600753 | 0.542 | 0.246 | DYX1C1 | Scerri et al.^[58]^ | 0.01057 | 0.9562 |
| rs807724 |  | 1.00 | 0.056 | DCDC2 | Schumacher et al^[57]^ | 0.2459 | 0.4844 |
| rs807701 |  | 1.00 | 0.246 | DCDC2 | Schumacher et al^[57]^ | 0.281 | 0.1365 |
| rs2143340 |  | 1.00 | 0.171 | KIAA0319 | Francks et al.^[54]^ | 0.3105 | 0.1583 |
| rs16889506 |  | 1.00 | 0.167 | KIAA0319 | Zhao et al^[32]^ | 0.4017 | 0.06928 |
| rs699463 |  | 1.00 | 0.14 | KIAA0319 | Zhao et al^[32]^ | -0.1253 | 0.5974 |
| rs9366577 |  | 1.00 | 0.052 | KIAA0319 | Zhao et al^[32]^ | -0.4468 | 0.2271 |
| rs793862 |  | 1.00 | 0.389 | DCDC2 | Schumacher et al^[57]^ | -0.1637 | 0.3257 |
| rs714939 |  | 1.00 | 0.398 | MRPL19 | Anthoni et al.^[55]^ | -0.01009 | 0.9515 |
| rs1091031 |  | 1.00 | 0.213 | KIAA0319 | Zhao et al^[32]^ | 0.2161 | 0.2814 |
| rs6732511 |  | 1.00 | 0.072 | MRPL19 | Anthoni et al.^[55]^ | 0.3474 | 0.2703 |
| rs2760157 |  | 1.00 | 0.438 | KIAA0319 | Zhao et al^[32]^ | -0.2325 | 0.1504 |
| rs6963842 |  | 1.00 | 0.209 | LAMB1 | Truong et al^[21]^ | 0.07187 | 0.722 |
| rs4571421 |  | 1.00 | 0.427 | LINC02118 | Gialluisi et al | -0.06395 | 0.6976 |

| LeadingSNPs | SNP | R^2^ | MAF | GENE | Identified by | RANdigit_BETA | P |
| --- | --- | --- | --- | --- | --- | --- | --- |
| rs3756821 |  |  | 0.288 | KIAA0319 | Zhao et al^[32]^ | 0.1813 | 0.1774 |
|  | rs3212236 | .493 | 0.451 | KIAA0319 | Harold et al^[52]^ | 0.09403 | 0.4411 |
|  | rs16889556 | .454 | 0.157 | KIAA0319 | Zhao et al^[32]^ | 0.3138 | 0.06367 |
|  | rs6935076 | .721 | 0.228 | KIAA0319 | Cope et al^[53]^ | 0.2665 | 0.0686 |
| rs761100 |  |  | 0.171 | KIAA0319 | Harold et al^[52]^ | -0.1375 | 0.4036 |
|  | rs2038137 | 0.927 | 0.161 | KIAA0319 | Zhao et al^[32]^ | -0.06458 | 0.703 |
|  | rs4504469 | 0.626 | 0.175 | KIAA0319 | Francks et al.^[54]^ | -0.04529 | 0.7787 |
|  | rs2179515 | 0.914 | 0.16 | KIAA0319 | Harold et al^[52]^ | -0.07904 | 0.6394 |
| rs807507 |  |  | 0.246 | KIAA0319 | Zhao et al^[32]^ | -0.2378 | 0.0915 |
|  | rs12193738 | 0.971 | 0.246 | KIAA0319 | Zhao et al^[32]^ | -0.3078 | 0.02874 |
|  | rs3903801 | 0.858 | 0.257 | KIAA0319 | Zhao et al^[32]^ | -0.367 | 0.007596 |
| rs1000585 |  |  | 0.422 | MRPL19 | Anthoni et al.^[55]^ | 0.07233 | 0.5546 |
|  | rs917235 | 0.944 | 0.424 | MRPL19 | Anthoni et al.^[55]^ | 0.02324 | 0.8503 |
| rs3743204 |  |  | 0.16 | DYX1C1 | Wigg et al.^[56]^ | -0.2702 | 0.1103 |
|  | rs600753 | 0.542 | 0.246 | DYX1C1 | Scerri et al.^[58]^ | -0.3075 | 0.03275 |
| rs807724 |  | 1.00 | 0.056 | DCDC2 | Schumacher et al^[57]^ | 0.1479 | 0.5838 |
| rs807701 |  | 1.00 | 0.246 | DCDC2 | Schumacher et al^[57]^ | -0.09981 | 0.4872 |
| rs2143340 |  | 1.00 | 0.171 | KIAA0319 | Francks et al.^[54]^ | 0.03469 | 0.834 |
| rs16889506 |  | 1.00 | 0.167 | KIAA0319 | Zhao et al^[32]^ | -0.2964 | 0.07528 |
| rs699463 |  | 1.00 | 0.14 | KIAA0319 | Zhao et al^[32]^ | -0.005639 | 0.9747 |
| rs9366577 |  | 1.00 | 0.052 | KIAA0319 | Zhao et al^[32]^ | 0.1521 | 0.5828 |
| rs793862 |  | 1.00 | 0.389 | DCDC2 | Schumacher et al^[57]^ | 0.1962 | 0.1192 |
| rs714939 |  | 1.00 | 0.398 | MRPL19 | Anthoni et al.^[55]^ | 0.08321 | 0.5073 |
| rs1091031 |  | 1.00 | 0.213 | KIAA0319 | Zhao et al^[32]^ | -0.02869 | 0.851 |
| rs6732511 |  | 1.00 | 0.072 | MRPL19 | Anthoni et al.^[55]^ | -0.1502 | 0.521 |
| rs2760157 |  | 1.00 | 0.438 | KIAA0319 | Zhao et al^[32]^ | 0.3217 | 0.007647 |
| rs6963842 |  | 1.00 | 0.209 | LAMB1 | Truong et al^[21]^ | -0.1296 | 0.3962 |
| rs4571421 |  | 1.00 | 0.427 | LINC02118 | Gialluisi et al | 0.07006 | 0.5709 |

| LeadingSNPs | SNP | R^2^ | MAF | GENE | Identified by | RANdice_BETA | P |
| --- | --- | --- | --- | --- | --- | --- | --- |
| rs3756821 |  |  | 0.288 | KIAA0319 | Zhao et al^[32]^ | 0.01742 | 0.9306 |
|  | rs3212236 | .493 | 0.451 | KIAA0319 | Harold et al^[52]^ | -0.1308 | 0.47 |
|  | rs16889556 | .454 | 0.157 | KIAA0319 | Zhao et al^[32]^ | 0.3309 | 0.1894 |
|  | rs6935076 | .721 | 0.228 | KIAA0319 | Cope et al^[53]^ | 0.2217 | 0.3093 |
| rs761100 |  |  | 0.171 | KIAA0319 | Harold et al^[52]^ | -0.3239 | 0.1814 |
|  | rs2038137 | 0.927 | 0.161 | KIAA0319 | Zhao et al^[32]^ | -0.2705 | 0.2786 |
|  | rs4504469 | 0.626 | 0.175 | KIAA0319 | Francks et al.^[54]^ | -0.04529 | 0.7787 |
|  | rs2179515 | 0.914 | 0.16 | KIAA0319 | Harold et al^[52]^ | -0.07904 | 0.6394 |
| rs807507 |  |  | 0.246 | KIAA0319 | Zhao et al^[32]^ | -0.3752 | 0.07188 |
|  | rs12193738 | 0.971 | 0.246 | KIAA0319 | Zhao et al^[32]^ | -0.4382 | 0.0353 |
|  | rs3903801 | 0.858 | 0.257 | KIAA0319 | Zhao et al^[32]^ | -0.3638 | -0.3638 |
| rs1000585 |  |  | 0.422 | MRPL19 | Anthoni et al.^[55]^ | 0.2375 | 0.1897 |
|  | rs917235 | 0.944 | 0.424 | MRPL19 | Anthoni et al.^[55]^ | 0.1279 | 0.4825 |
| rs3743204 |  |  | 0.16 | DYX1C1 | Wigg et al.^[56]^ | -0.2238 | 0.3771 |
|  | rs600753 | 0.542 | 0.246 | DYX1C1 | Scerri et al.^[58]^ | -0.269 | 0.2102 |
| rs807724 |  | 1.00 | 0.056 | DCDC2 | Schumacher et al^[57]^ | 0.4428 | 0.2679 |
| rs807701 |  | 1.00 | 0.246 | DCDC2 | Schumacher et al^[57]^ | -0.07628 | 0.7199 |
| rs2143340 |  | 1.00 | 0.171 | KIAA0319 | Francks et al.^[54]^ | 0.1163 | 0.6358 |
| rs16889506 |  | 1.00 | 0.167 | KIAA0319 | Zhao et al^[32]^ | -0.6193 | 0.01216 |
| rs699463 |  | 1.00 | 0.14 | KIAA0319 | Zhao et al^[32]^ | 0.02178 | 0.9341 |
| rs9366577 |  | 1.00 | 0.052 | KIAA0319 | Zhao et al^[32]^ | -0.1243 | 0.7609 |
| rs793862 |  | 1.00 | 0.389 | DCDC2 | Schumacher et al^[57]^ | 0.3676 | 0.04883 |
| rs714939 |  | 1.00 | 0.398 | MRPL19 | Anthoni et al.^[55]^ | -0.5211 | 0.02064 |
| rs1091031 |  | 1.00 | 0.213 | KIAA0319 | Zhao et al^[32]^ | -0.2322 | 0.3027 |
| rs6732511 |  | 1.00 | 0.072 | MRPL19 | Anthoni et al.^[55]^ | -0.1502 | 0.7586 |
| rs2760157 |  | 1.00 | 0.438 | KIAA0319 | Zhao et al^[32]^ | 0.3571 | 0.04497 |
| rs6963842 |  | 1.00 | 0.209 | LAMB1 | Truong et al^[21]^ | -0.1296 | 0.3962 |
| rs4571421 |  | 1.00 | 0.427 | LINC02118 | Gialluisi et al | 0.1878 | 0.3039 |

| LeadingSNPs | SNP | R^2^ | MAF | GENE | Identified by | RANpicture_BETA | P |
| --- | --- | --- | --- | --- | --- | --- | --- |
| rs3756821 |  |  | 0.288 | KIAA0319 | Zhao et al^[32]^ | -0.2275 | 0.3634 |
|  | rs3212236 | .493 | 0.451 | KIAA0319 | Harold et al^[52]^ | -0.309 | 0.1737 |
|  | rs16889556 | .454 | 0.157 | KIAA0319 | Zhao et al^[32]^ | 0.07251 | 0.8179 |
|  | rs6935076 | .721 | 0.228 | KIAA0319 | Cope et al^[53]^ | -0.03168 | 0.9076 |
| rs761100 |  |  | 0.171 | KIAA0319 | Harold et al^[52]^ | -0.3531 | 0.2463 |
|  | rs2038137 | 0.927 | 0.161 | KIAA0319 | Zhao et al^[32]^ | -0.2208 | 0.4808 |
|  | rs4504469 | 0.626 | 0.175 | KIAA0319 | Francks et al.^[54]^ | 0.2975 | 0.3208 |
|  | rs2179515 | 0.914 | 0.16 | KIAA0319 | Harold et al^[52]^ | -0.2613 | 0.4032 |
| rs807507 |  |  | 0.246 | KIAA0319 | Zhao et al^[32]^ | -0.4958 | 0.05834 |
|  | rs12193738 | 0.971 | 0.246 | KIAA0319 | Zhao et al^[32]^ | -0.6397 | 0.01441 |
|  | rs3903801 | 0.858 | 0.257 | KIAA0319 | Zhao et al^[32]^ | -0.5604 | 0.02837 |
| rs1000585 |  |  | 0.422 | MRPL19 | Anthoni et al.^[55]^ | 0.1688 | 0.4569 |
|  | rs917235 | 0.944 | 0.424 | MRPL19 | Anthoni et al.^[55]^ | 0.06339 | 0.7814 |
| rs3743204 |  |  | 0.16 | DYX1C1 | Wigg et al.^[56]^ | -0.1855 | 0.5596 |
|  | rs600753 | 0.542 | 0.246 | DYX1C1 | Scerri et al.^[58]^ | 0.03566 | 0.8947 |
| rs807724 |  | 1.00 | 0.056 | DCDC2 | Schumacher et al^[57]^ | 0.2225 | 0.6565 |
| rs807701 |  | 1.00 | 0.246 | DCDC2 | Schumacher et al^[57]^ | 0.1338 | 0.616 |
| rs2143340 |  | 1.00 | 0.171 | KIAA0319 | Francks et al.^[54]^ | -0.03446 | 0.9109 |
| rs16889506 |  | 1.00 | 0.167 | KIAA0319 | Zhao et al^[32]^ | -0.5959 | 0.0555 |
| rs699463 |  | 1.00 | 0.14 | KIAA0319 | Zhao et al^[32]^ | -0.05508 | 0.8675 |
| rs9366577 |  | 1.00 | 0.052 | KIAA0319 | Zhao et al^[32]^ | 0.006441 | 0.99 |
| rs793862 |  | 1.00 | 0.389 | DCDC2 | Schumacher et al^[57]^ | 0.1134 | 0.6286 |
| rs714939 |  | 1.00 | 0.398 | MRPL19 | Anthoni et al.^[55]^ | 0.06339 | 0.7814 |
| rs1091031 |  | 1.00 | 0.213 | KIAA0319 | Zhao et al^[32]^ | -0.102 | 0.7185 |
| rs6732511 |  | 1.00 | 0.072 | MRPL19 | Anthoni et al.^[55]^ | 0.4346 | 0.3207 |
| rs2760157 |  |  |  | KIAA0319 | Zhao et al^[32]^ | 0.6002 | 0.00714 |
| rs6963842 |  | 1.00 | 0.209 | LAMB1 | Truong et al^[21]^ | -0.6731 | 0.01694 |
| rs4571421 |  | 1.00 | 0.427 | LINC02118 | Gialluisi et al | 0.5008 | 0.02859 |

| LeadingSNPs | SNP | R^2^ | MAF | GENE | Identified by | RANcolor_BETA | P |
| --- | --- | --- | --- | --- | --- | --- | --- |
| rs3756821 |  |  | 0.288 | KIAA0319 | Zhao et al^[32]^ | -0.02602 | 0.9336 |
|  | rs3212236 | .493 | 0.451 | KIAA0319 | Harold et al^[52]^ | -0.1553 | 0.584 |
|  | rs16889556 | .454 | 0.157 | KIAA0319 | Zhao et al^[32]^ | 0.5779 | 0.143 |
|  | rs6935076 | .721 | 0.228 | KIAA0319 | Cope et al^[53]^ | 0.5764 | 0.09113 |
| rs761100 |  |  | 0.171 | KIAA0319 | Harold et al^[52]^ | -0.4339 | 0.2545 |
|  | rs2038137 | 0.927 | 0.161 | KIAA0319 | Zhao et al^[32]^ | -0.2171 | 0.579 |
|  | rs4504469 | 0.626 | 0.175 | KIAA0319 | Francks et al.^[54]^ | 0.03147 | 0.9329 |
|  | rs2179515 | 0.914 | 0.16 | KIAA0319 | Harold et al^[52]^ | -0.3372 | 0.3878 |
| rs807507 |  |  | 0.246 | KIAA0319 | Zhao et al^[32]^ | -0.7677 | 0.01853 |
|  | rs12193738 | 0.971 | 0.246 | KIAA0319 | Zhao et al^[32]^ | -0.9117 | 0.005088 |
|  | rs3903801 | 0.858 | 0.257 | KIAA0319 | Zhao et al^[32]^ | -0.7646 | 0.01652 |
| rs1000585 |  |  | 0.422 | MRPL19 | Anthoni et al.^[55]^ | -0.01906 | 0.9463 |
|  | rs917235 | 0.944 | 0.424 | MRPL19 | Anthoni et al.^[55]^ | -0.09718 | 0.733 |
| rs3743204 |  |  | 0.16 | DYX1C1 | Wigg et al.^[56]^ | -0.3934 | 0.3213 |
|  | rs600753 | 0.542 | 0.246 | DYX1C1 | Scerri et al.^[58]^ | -0.0625 | 0.8523 |
| rs807724 |  | 1.00 | 0.056 | DCDC2 | Schumacher et al^[57]^ | 0.3628 | 0.5605 |
| rs807701 |  | 1.00 | 0.246 | DCDC2 | Schumacher et al^[57]^ | -0.02533 | 0.9395 |
| rs2143340 |  | 1.00 | 0.171 | KIAA0319 | Francks et al.^[54]^ | -0.155 | 0.6858 |
| rs16889506 |  | 1.00 | 0.167 | KIAA0319 | Zhao et al^[32]^ | -1.391 | 3.168e-4 |
| rs699463 |  | 1.00 | 0.14 | KIAA0319 | Zhao et al^[32]^ | 0.126 | 0.7591 |
| rs9366577 |  | 1.00 | 0.052 | KIAA0319 | Zhao et al^[32]^ | 0.7692 | 0.2296 |
| rs793862 |  | 1.00 | 0.389 | DCDC2 | Schumacher et al^[57]^ | 0.02151 | 0.9412 |
| rs714939 |  | 1.00 | 0.398 | MRPL19 | Anthoni et al.^[55]^ | -0.03206 | 0.9126 |
| rs1091031 |  | 1.00 | 0.213 | KIAA0319 | Zhao et al^[32]^ | -0.2058 | 0.5587 |
| rs6732511 |  | 1.00 | 0.072 | MRPL19 | Anthoni et al.^[55]^ | 0.3571 | 0.5124 |
| rs2760157 |  | 1.00 | 0.438 | KIAA0319 | Zhao et al^[32]^ | 1.103 | 6.967e-05 |
| rs6963842 |  | 1.00 | 0.209 | LAMB1 | Truong et al^[21]^ | -0.2514 | 0.4761 |
| rs4571421 |  | 1.00 | 0.427 | LINC02118 | Gialluisi et al | 0.1689 | 0.5537 |
